# Exposure to a mixture of BMAA and MCLR synergistically modulates behavior in larval zebrafish while exacerbating molecular changes related to neurodegeneration

**DOI:** 10.1101/2020.07.15.205617

**Authors:** Rubia M. Martin, Michael S. Bereman, Kurt C. Marsden

**Affiliations:** Department of Biological Sciences, North Carolina State University, Raleigh, NC

**Keywords:** Cyanotoxins, Mixtures, Synergism, Zebrafish, Behavior, Proteomics

## Abstract

Exposure to toxins produced by cyanobacteria (i.e., cyanotoxins) is an emerging health concern due to their increased occurrence and previous associations with neurodegenerative disease including amyotrophic lateral sclerosis (ALS). The objective of this study was to evaluate the neurotoxic effects of a mixture of two co-occurring cyanotoxins, β-methylamino-L-alanine (BMAA) and microcystin leucine and arginine (MCLR), using the larval zebrafish model. We combined high-throughput behavior based toxicity assays with discovery proteomic techniques to identify behavioral and molecular changes following 6 days of exposure. While neither toxin caused mortality, morphological defects, or altered general locomotor behavior in zebrafish larvae, both toxins increased acoustic startle sensitivity in a dose-dependent manner by at least 40% (p<0.0001). Furthermore, startle sensitivity was enhanced by an additional 40% in larvae exposed to the BMAA/MCLR mixture relative to those exposed to the individual toxins. Supporting these behavioral results, our proteomic analysis revealed a 4-fold increase in the number of differentially expressed proteins (DEPs) in the mixture exposed group. Additionally, prediction analysis reveals activation and/or inhibition of 8 enriched canonical pathways (enrichment p-value<0.01; z-score≥|2|), including ILK, Rho Family GTPase, RhoGDI, and calcium signaling pathways, which have been implicated in neurodegeneration. We also found that expression of TDP-43, of which cytoplasmic aggregates are a hallmark of ALS pathology, was significantly upregulated by 5.7-fold following BMAA/MCLR mixture exposure. Together, our results emphasize the importance of including mixtures of cyanotoxins when investigating the link between environmental cyanotoxins and neurodegeneration as we reveal that BMAA and MCLR interact *in vivo* to enhance neurotoxicity.

## 1. Introduction

Amyotrophic lateral sclerosis (ALS) is the most common neurodegenerative disease of midlife and is rapidly fatal with a median survival period of three years from symptom onset (Brown and Al-Chalabi 2017). It is defined by a progressive loss of both upper and lower motor neurons, resulting in muscle spasticity, weakness, and atrophy (Swinnen and Robberecht 2014). Approximately 10% of ALS cases are classified as familial due to the inheritance of single gene mutations (Renton et al. 2014). For the remaining 90% of cases the disease etiology is unknown and likely stems from a complex interplay between genetic and environmental factors. (Ingre et al. 2015; Jones 2009). Although the contribution of environmental factors to sporadic ALS (sALS) is difficult to assess as the search space is infinite, several studies have associated ALS incidence with exposure to heavy metals, pesticides, and electromagnetic fields (reviewed in (Bozzoni 2016)). In addition, there is also strong evidence that exposure to cyanotoxins is a major risk factor for sALS (Bradley and Mash 2009).

The link between beta-methylamino-L-alanine (BMAA), a toxin produced by a diverse taxa of cyanobacteria (Cox et al. 2005) and sALS was first observed on the island of Guam in the 1950s (Kurland and Mulder 1955). The indigenous population of Guam succumbed to an ALS/parkinsonism-dementia (PD) neurodegenerative complex with a 100-fold greater incidence than the rest of the world (Bradley and Mash 2009). The elevated rates of ALS/PD in Guam were attributed to BMAA exposure as the indigenous population consumed flour made from BMAA-containing cycad seeds as well as flying foxes in which BMAA was biomagnified up to 10,000-fold greater than in free living bacteria (3,556 μg/g BMAA) (Banack and Cox 2003). Since then, numerous studies have implicated BMAA in sALS cases outside of Guam, including clusters of ALS along the French Mediterranean coast, New Hampshire, and Maryland (Caller et al. 2009; Field et al. 2013; Masseret et al. 2013). These epidemiological studies provide evidence that exposure to BMAA is associated with neurodegeneration. Epidemiological findings are further supported by laboratory studies in which BMAA was found to cause neurotoxic effects consistent with neurodegenerative disease (Beri et al. 2017; Karlsson et al. 2017). Furthermore, neonatal BMAA exposure in rats has been shown to produce motor defects in rats (Scott et al. 2017), indicating that exposure during neural developmental may enhance sALS risk. However, a major limitation for these and many other BMAA studies is that BMAA is just one of many toxic metabolites produced by cyanobacteria, some of which have been reported to co-occur with BMAA around the world (Banack et al. 2015; Sabart et al. 2015). Thus, to obtain a more thorough understanding of the risk posed by exposure to cyanotoxic blooms, it is essential to investigate the toxicity of other cyanotoxins with BMAA.

BMAA at low concentrations (~10 μM) in combination with other non-cyanotoxic neurotoxins has been found to potentiate neuronal damage *in vitro* (Lobner et al. 2006). More recently, our laboratory demonstrated that co-exposure to BMAA and its isomers (i.e., AEG and 2,4DAB) at low concentrations (~166 μM) produces a synergistic interaction *in vitro*, perturbing regulation of various canonical pathways, bioprocesses, and upstream regulators involved in neurodegenerative processes (Martin et al. 2019). Recent studies have shown that microcystin leucine-arginine (MCLR) is also a potent neurotoxin (Tzima et al. 2017; Wu et al. 2016), and like BMAA, MCLR can bioaccumulate in tissues (Wang et al. 2008; Zhao et al. 2015). Microcystins are the most abundant cyanotoxins in the environment and have been shown to co-occur with BMAA (Banack et al. 2015; Jungblut et al. 2018; Metcalf et al. 2012). Therefore, co-exposure to BMAA and MCLR is of increasing toxicological significance.

To identify potential neurotoxic effects *in vivo*, we exposed larval zebrafish to BMAA and/or MCLR and assessed neural function with a set of behavioral assays using a high-throughput testing platform. While neither BMAA nor MCLR caused changes in locomotion, both toxins increased acoustic startle sensitivity in a dose-dependent manner. Furthermore, a mixture of BMAA and MCLR enhanced toxicity in the startle assay. Finally, we examined the protein profile of larval zebrafish exposed to the BMAA/MCLR mixture and identified molecular signatures consistent with neurodegeneration, including upregulation of the ALS-associated protein TDP-43 (Mackenzie et al. 2010). Together, our data highlight the importance of studying toxic mixtures and reveal novel mechanisms that may link cyanotoxin exposure to sALS.

## 2. Materials and Methods

### 2.1. Chemicals

Synthetic BMAA standards were obtained from Sigma Aldrich (St. Louis, MO, USA), and purified MCLR (purity > 95%) was obtained from Enzo Life Sciences, Inc. (Farmingdale, NY, USA). Water, acetonitrile, methanol, acetic acid, and formic acid were all Optima LC–MS grade solvents purchased from Fisher Scientific (Tewksbury, MA, USA). A stock solution of BMAA at 10 mg.mL^−1^ and MCLR at 1 mg.mL^−1^ was used for all dilutions. All BMAA dilutions were prepared in HPLC grade water while MCLR dilutions were prepared in DMSO.

### 2.2. Zebrafish husbandry and exposures

All animal use and procedures were approved by the North Carolina State University IACUC. Zebrafish (*Danio rerio*) embryos from multiple crosses of wild-type tupfel longfin (TLF) strain adults were collected and placed into Petri dishes containing E3 medium, and unfertilized eggs were removed as described previously (Burgess and Granato 2007). Embryos from all clutches were mixed and randomly sorted into 24 well plates (8-10 animals per well) containing 1 mL of E3 per well.

At 6 hpf, all E3 was removed and replaced with vehicle (HPLC-grade water),100, 500, or 1000 μM BMAA in E3, vehicle (DMSO), 1, 2.5, 5, or 10 μM MCLR in E3, or 100 μM BMAA plus 1 μM MCLR in E3. All treatments were performed in triplicate and were repeated in each of 3 separate experiments. Embryos were incubated at 29°C on a 14h:10h light-dark cycle, and 100% of the media was exchanged for fresh solutions daily. Embryos/ larvae were exposed to treatments until 6 days post fertilization (6 dpf).

### 2.3. Behavior assays and analysis

All 6 dpf larvae were thoroughly screened for developmental defects, and those with uninflated swim bladder, edema, or other morphological defects were removed from analysis. Screened larvae were adapted to the testing lighting and temperature conditions for 30 minutes prior to testing. Behavior testing was done as previously described (Burgess and Granato 2007; Marsden et al. 2018). Briefly, 6 dpf larvae were transferred to individual 9 mm round wells on a 36-well laser-cut acrylic testing grid. Larvae acclimated for 5 min and then spontaneous locomotor activity was recorded for 18.5 min at 640 × 640 px resolution at 50 frames per sec (fps) using a Photron mini UX-50 high-speed camera. The same set of larvae were then presented with a total of 60 acoustic stimuli, 10 at each of 6 intensities (13.6, 25.7, 29.2, 35.5, 39.6, and 53.6 dB), with a 20s interstimulus interval (ISI). Startle responses were recorded at 1000 fps. Stimuli were delivered by an acoustic-vibrational shaker (Bruel and Kjaer) to which the testing grid was directly mounted. All stimuli were calibrated with a PCB Piezotronics accelerometer (#355B04) and signal conditioner (#482A21), and voltage outputs were converted to dB using the formula dB = 20 log V. Analysis of recorded behaviors was done using FLOTE software as described previously (Burgess and Granato 2007; Marsden et al. 2018). Short-latency C-bends (SLCs) and long-latency C-bends (LLCs) were determined by defined kinematic parameters. A startle sensitivity index was calculated for individual larvae by calculating the area under the curve of startle frequency versus stimulus intensity using Prism 8 software (GraphPad). Statistical analyses were performed using JMP pro 14 from SAS institute, Cary, NC. Data were analyzed for effects between the groups (comparison of means), using Tukey-Kramer HSD, Alpha 0.05. Violin plots were generated using Prism 8.

### 2.4 Proteomics analysis

#### Sample preparation and LC MS/MS

Details of sample preparation, protein extraction and digestion via filter aided sample preparation (FASP) can be found in Supplemental Methods. Details regarding the LC-MS/MS data collection are also provided in the Supplemental Methods. Raw data files obtained in this experiment have been made available on the Chorus LC-MS data repository and can be assessed under the project ID#1679.

#### Proteomics Data Analysis

Details for the label free quantitation (LFQ) have been previously described here (Martin et al. 2019). In brief, LFQ was performed in MaxQuant (version.1.5.60), and data were searched against the *Danio rerio* Swiss Prot protein database (# protein sequences = 56 281, accessed 03/22/2019). Comparison of LFQ intensities across the whole set of measurements was investigated using Perseus software (version 1.5.1.6), where calculation of statistical significance was determined using two-way Student-t test and FPR (p ≤0.05).

#### Pathway Analysis

Ingenuity Pathway Analysis (IPA) software was used to identify the function, specific processes, and enriched pathways of the differentially expressed proteins using the “Core Analysis” function. Only significantly differentially expressed proteins (p ≤ 0.05) were submitted to IPA. We used an empirical background protein database to evaluate the significance of pathway enrichment. The database was created by using all of the proteins that were detected in our samples (Bereman et al. 2018; Khatri and Drăghici 2005).

## 3. Results

An overview of the experimental design is illustrated in **Figure 1**. In brief, we first conducted a dose-response study to determine the no observed adverse effect levels (NOAELs) to be implemented in subsequent mixture analyses. Zebrafish larvae were exposed to increasing concentrations of BMAA or MCLR from 6 hours post-fertilization (hpf) to 6 days post-fertilization (dpf). At 6 dpf, neurotoxicity was evaluated via two behavioral assays: spontaneous movement and acoustic startle response assays. Based on these data, a mixture was created using BMAA and MCLR at their respective NOAELs in which zebrafish larvae were exposed as before, followed by behavior analysis to identify potential interactions between BMAA and MCLR. Finally, to investigate perturbed molecular pathways associated with cyanotoxic mixture exposure, the mixture-exposed group and their respective controls were subjected to shotgun proteomics.

**Fig. 1.**
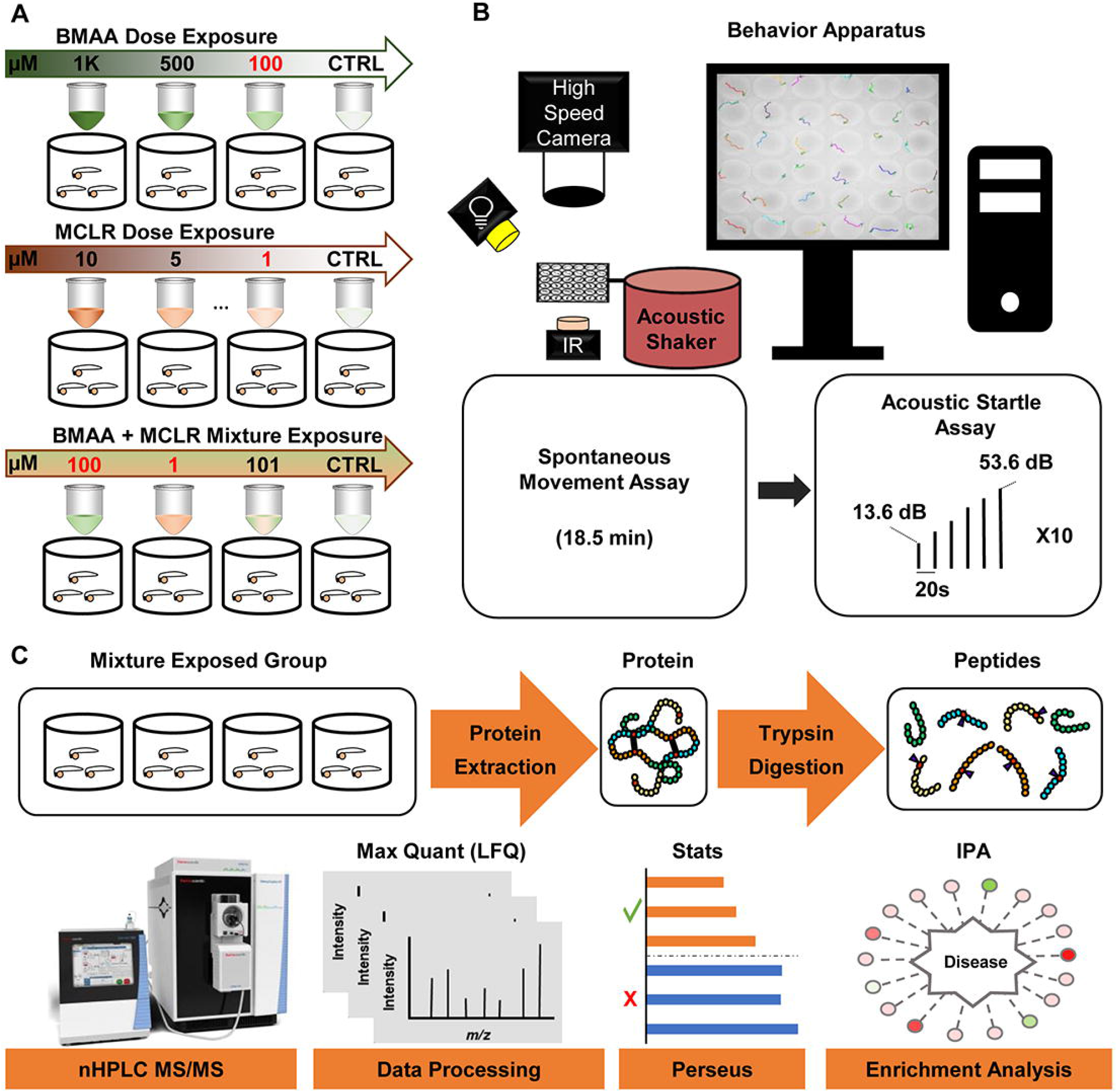
Experimental Design. (A) Cyanotoxin exposure plan for zebrafish from 6 hpf to 6 dpf. (B) High throughput behavior testing apparatus: multi-well testing grid is mounted on an acoustic shaker above an infrared (IR) LED array and below an IR-sensitive high-speed camera. A white LED is mounted above the grid to simulate daylight conditions. Videos are analyzed with automated tracking software (FLOTE). (C) Proteomics workflow: zebrafish larvae from the mixture exposed groups were pooled for protein extraction and tryptic digestion of extracted proteins into peptides. nLC-MS/MS label free protein quantitation via MaxQuant statistical analysis via Perseus software and enrichment analysis via ingenuity pathway analysis (IPA).

### 3.1 BMAA and MCLR Dose Response Study: identification of NOAELs

To determine if exposure to environmentally relevant concentrations of BMAA or MCLR cause neurotoxicity in wild-type zebrafish, we treated TLF strain embryos from 6 hpf to 6 dpf with increasing concentrations of BMAA (100, 500, and 1000 μM) and MCLR (1, 2.5, 5, and 10 μM). We did not observe increased mortality, morbidity, or any overt developmental phenotypes in any of the exposed groups of larvae. First, we examined the effect of BMAA and MCLR on general locomotion (**Figure 2**) using a custom built, high-throughput behavior platform and unbiased, automated FLOTE tracking and analysis software (Burgess and Granato 2007). In order to investigate if various concentrations of BMAA and/or MCLR could alter spontaneous movement, 6 dpf larvae were adapted to the testing conditions for 30 min, transferred to a multi-well grid mounted below a high-speed camera, habituated for 5 additional minutes, and then their spontaneous movements were recorded for 18.5 min. We detected no significant differences in total distance travelled for zebrafish larvae treated with either BMAA or MCLR compared to their respective vehicle controls (**Figure 2A**). Average speed was also unchanged in all groups, except for larvae treated with 1000 μM BMAA, whose speed was significantly reduced (**Figure 2B**). We also examined the frequency of turning and swimming behaviors, as defined by specific kinematic parameters (Hao le et al. 2012). There were no significant differences in the ratio of turns to swims in BMAA or MCLR treated larvae (**Figure 2C**). The overall frequency of these movements was also unchanged, except for a slight increase in turn frequency in 10 μM MCLR-treated larvae (**Supplemental Figure 1**). These data indicate that developmental exposure to BMAA or MCLR does not substantially affect general locomotor activity in larval zebrafish.

**Fig. 2.**
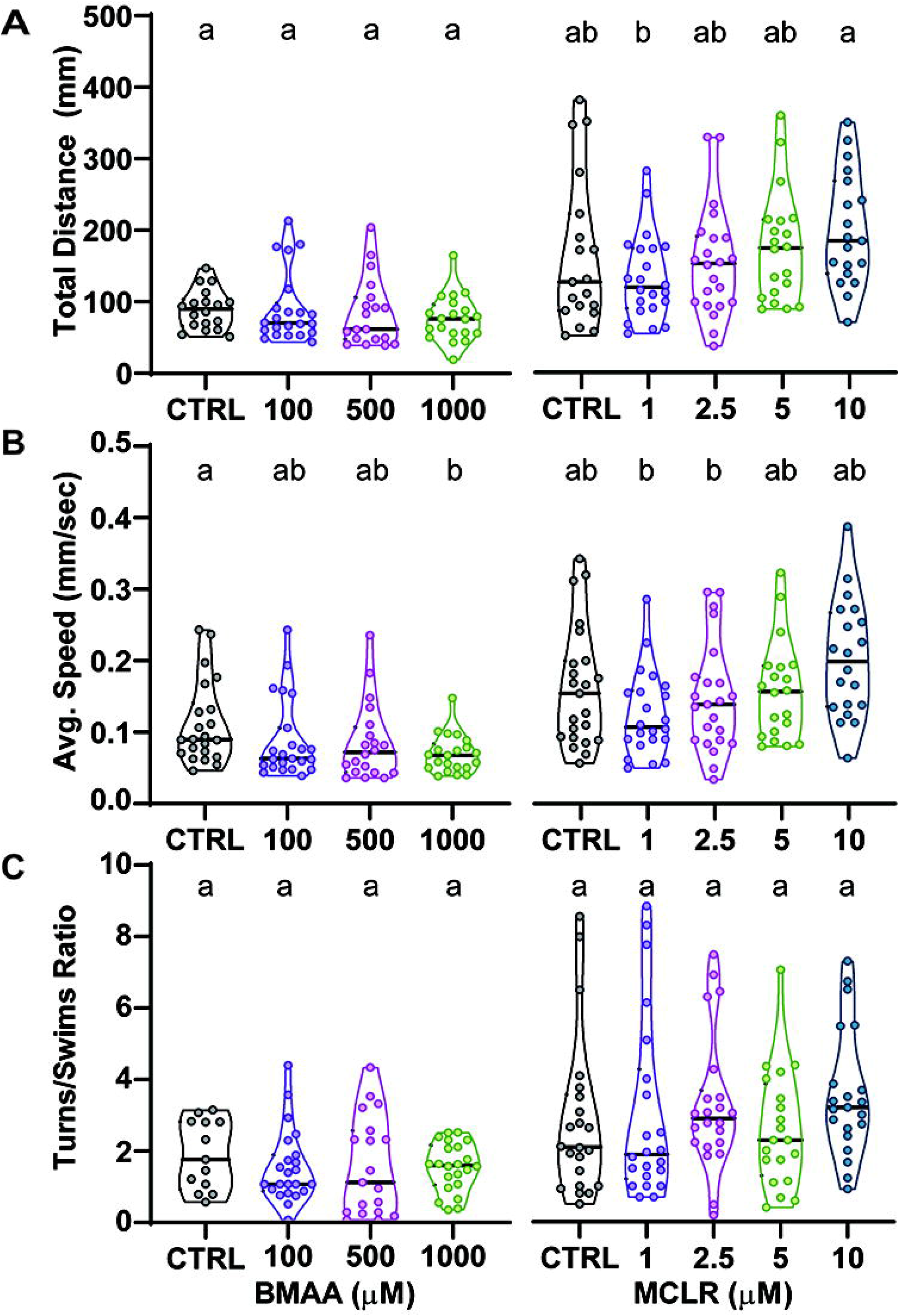
BMAA and MCLR do not substantially alter general locomotor activity. (A) Violin plots depict the total distance travelled during the 18.5 min spontaneous movement assay for each larva. (B) Average speed across the same assay. (C) The ratio of turning movements to swimming movements performed during the spontaneous movement assay. Levels not connected by the same letter are significantly different– Tukey-Kramer HSD, Alpha 0.05.

We then we examined sensorimotor function using an acoustic startle assay consisting of 60 total stimuli, 10 at each of 6 intensities with a 20 sec inter-stimulus interval (**Figure 3**). In response to an acoustic stimulus, zebrafish larvae perform one of two types of high-velocity startle behaviors: Short-latency C-bends (SLCs), which rely on the Mauthner neurons (Burgess and Granato 2007), or Long-latency C-bends (LLCs), which are independent of the Mauthner cells but require a set of prepontine neurons (Marquart et al. 2019). To investigate if BMAA or MCLR alters startle performance, we measured SLC and LLC frequency across the 60-stimulus assay (**Figure 3). Figure 3A** highlights both the SLC and LLC response frequency disparities between zebrafish larvae exposed to 1000 μM BMAA and vehicle control. 1000 μM BMAA increases SLC responses while decreasing LLC responses, indicating that BMAA shifts the behavioral response bias toward SLCs.

**Fig. 3.**
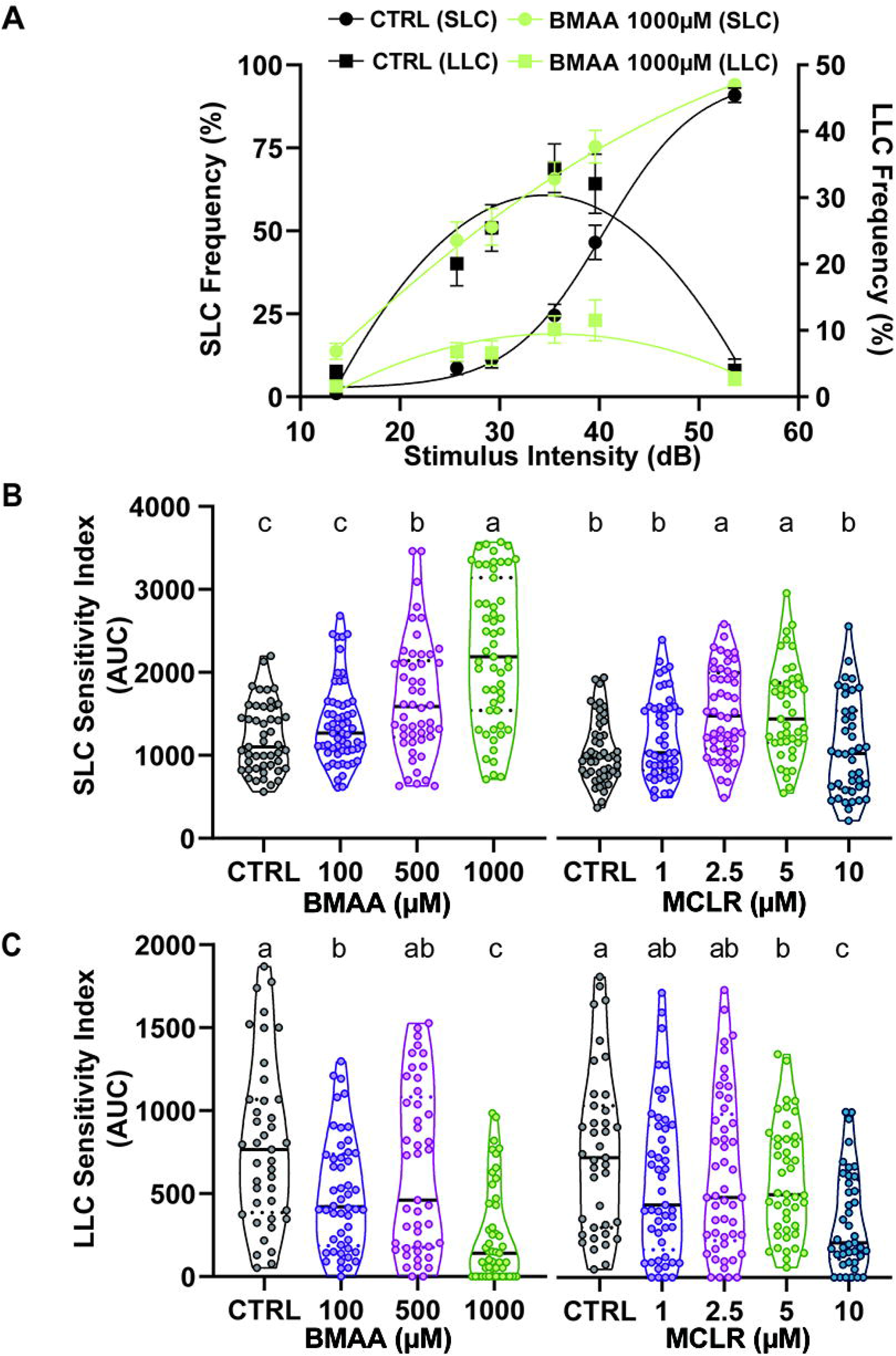
BMAA and MCLR significantly increase acoustic startle sensitivity. (A) Startle frequency curves for short-latency C-bends (SLCs, left axis, sigmoidal curve fits) and long-latency C-bends (LLCs, right axis, sigmoidal curve fits) in control larvae (black) and treated larvae (1000 μM BMAA; green). n = 54 larvae; mean ± SEM. (B) and (C), SLC and LLC sensitivity indices, calculated for each larva using the curves as in (A). Levels not connected by the same letter are significantly different–Tukey-Kramer HSD, Alpha 0.05.

To quantify SLC and LLC sensitivity, we calculated the area under the startle frequency curves in **Figure 3A** for each individual larva to create a startle sensitivity index (Marsden et al. 2018). Both BMAA and MCLR increased SLC sensitivity in a dose-dependent manner (**Figure 3B**). LLC responses decreased in a dose-dependent manner in both BMAA and MCLR-treated larvae, supporting an overall shift in response bias (**Figure 3C**). These data indicate that environmentally relevant concentrations of BMAA and MCLR enhance activity of the SLC circuit. In addition, these startle sensitivity data reveal NOAELs for BMAA (100 μM) and MCLR (1 μM), with NOAEL was defined as the highest non-statistically significant dose tested.

### 3.2. BMAA and MCLR Mixture Study: Interaction Amongst Cyanotoxins at a Behavioral Level

We next aimed to assess whether BMAA and MCLR interact *in vivo* by measuring the effects of a mixture of BMAA and MCLR at their respective NOAELs in larval zebrafish. We exposed wild-type zebrafish embryos from 6 hpf to 6 dpf to 4 different treatment conditions: 1) vehicle controls, 2) 100 μM BMAA, 3) 1 μM MCLR, and 4) 100 μM BMAA + 1 μM MCLR. As before, no overt developmental or morphological defects were observed in any exposed larvae. We then measured general locomotor activity and sensorimotor function using the same assays described above. In this cohort of animals, 100 μM BMAA very slightly decreased total distance traveled over 18.5 min **(Figure 4A),** and all three treatment groups showed a minor reduction in average speed during the assay (**Figure 4B**). No differences were observed in turning or swimming behaviors (**Figure 4C**). These data reinforce the results from our dose-response study that BMAA and MCLR do not substantially alter locomotor activity.

**Fig. 4.**
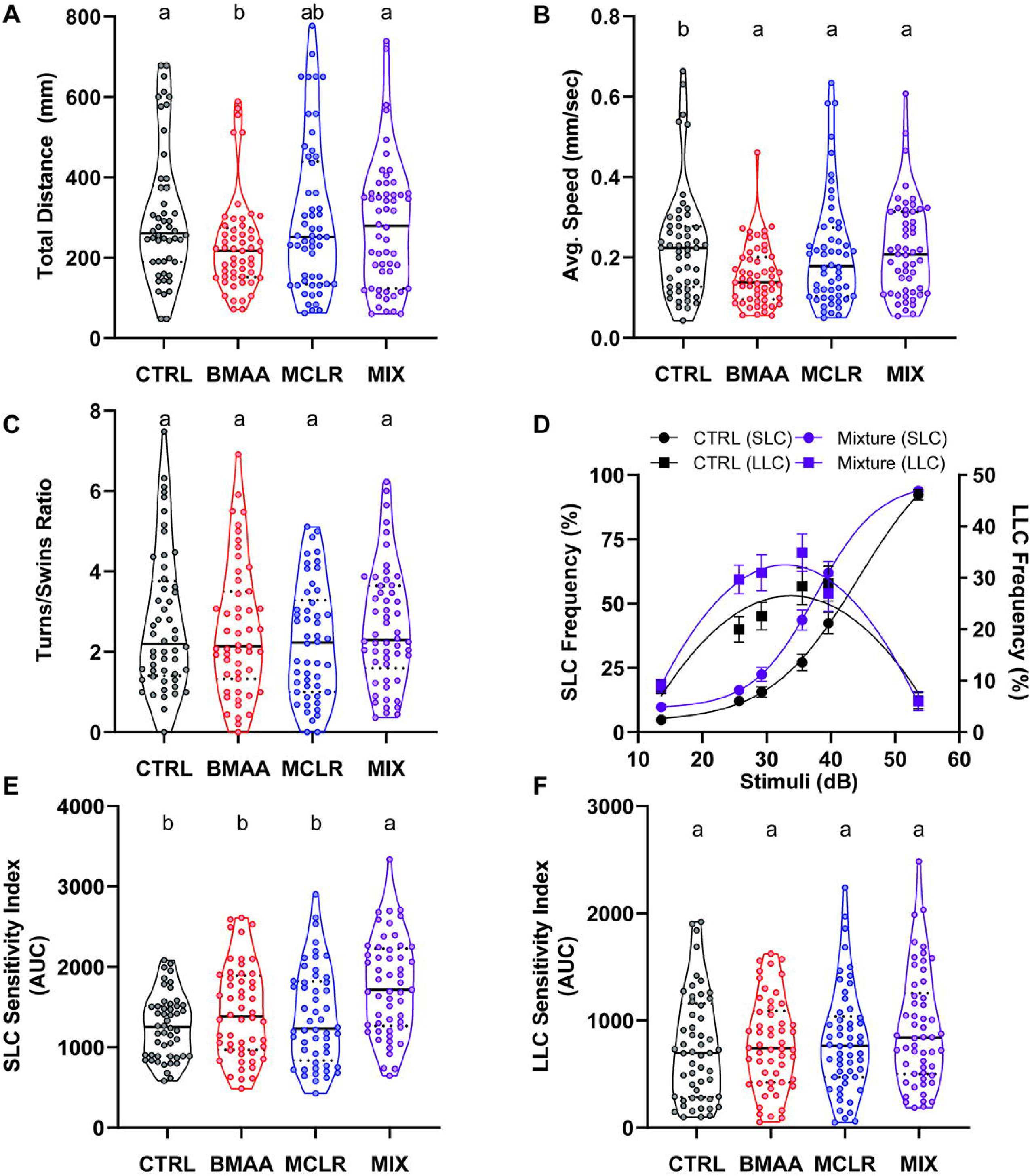
BMAA and MCLR interact to enhance startle sensitivity. (A) Violin plots depict the distribution of the total distance travelled during the 18.5 min spontaneous movement assay for each larva. (B) Violin plot of average speed. (C) Ratio of turns to swims. (D) Startle frequency curves for SLCs (left axis) and LLCs (right axis) in control larvae (0 μM of cyanotoxin, black) and mixture-treated larvae (100 μM BMAA plus 1 μM MCLR, purple). n = 54 siblings; mean ± SEM. (E,F) SLC and LLC indices for each larva. Levels not connected by the same letter are significantly different–Tukey-Kramer HSD, Alpha 0.05.

We next measured startle frequency in the same 4 groups of larvae. **Figure 4D** highlights both the SLC and LLC response frequency disparities between zebrafish larvae exposed to BMAA/MCLR mixture solution (101μM) and controls. Neither 100 μM BMAA nor 1 μM MCLR altered startle behavior, as both SLC (**Figure 4E**) and LLC sensitivity indices (**Figure 4F**) were unchanged. The 101 μM BMAA/MCLR mixture, however, significantly enhanced SLC sensitivity (**Figure 4E**) while leaving LLC sensitivity unchanged (**Figure 4F**), in contrast to the effect of BMAA alone (**Figure 3**). These data demonstrate not only that BMAA and MCLR interact *in vivo* to enhance SLC circuit activity, but because of the different effects of the mixture and the individual toxins on LLC responses (**Figure 3C vs. Figure 4F**), they also suggest that different cellular and/or molecular mechanisms are impacted by the mixture.

### 3.3 Global Proteomics Study: Interaction amongst cyanotoxins at a molecular level

To explore the molecular underpinnings associated with these behavioral phenotypes, we performed shotgun proteomics on larval zebrafish exposed to 100 μM BMAA and 1 μM MCLR alone and in combination. After behavioral testing, we carefully collected and flash-froze the treated zebrafish larvae, followed by protein extraction and digestion. Proteomes of larvae for each treatment condition were analyzed by LC-MS/MS, and approximately 3100 proteins were identified in each sample.

Differentially expressed proteins (DEPs) were determined by comparing the mean abundance within treatment to the control group for each protein using a two-way Student-t-test (P<0.05) (Tyanova et al. 2016). DEPs in all treatments can be found in **Supplemental Tables 2-4**. Volcano plots were used to visualize statistically significant differences in protein abundance across treatments in comparison to controls (**Supplemental Figure 2**). Notably, the BMAA/MCLR mixture induced the greatest molecular perturbation, with 259 DEPs compared to 79 for BMAA and 112 for MCLR, representing a 2.5-fold increase for the mixture-exposed group (**Figure 5A**). Although minimal overlap in DEPs between treatments was observed, we were intrigued by the nine proteins that were significantly differentially expressed in all three treatment groups (**Figure 5B; Supplemental table 1**). Out of these nine proteins, four proteins were mapped in the enrichment analysis (**Table 1**), and their general cellular functions include roles in cellular assembly, organization, and development. Interestingly, exposure to the BMAA/MCLR mixture also enhanced the abundance of these four DEPs by at least 2.5-fold relative to the individual cyanotoxins. These results reflect an enhanced toxicity after BMAA/MCLR mixture exposure *in vivo*.

**Table 1:**
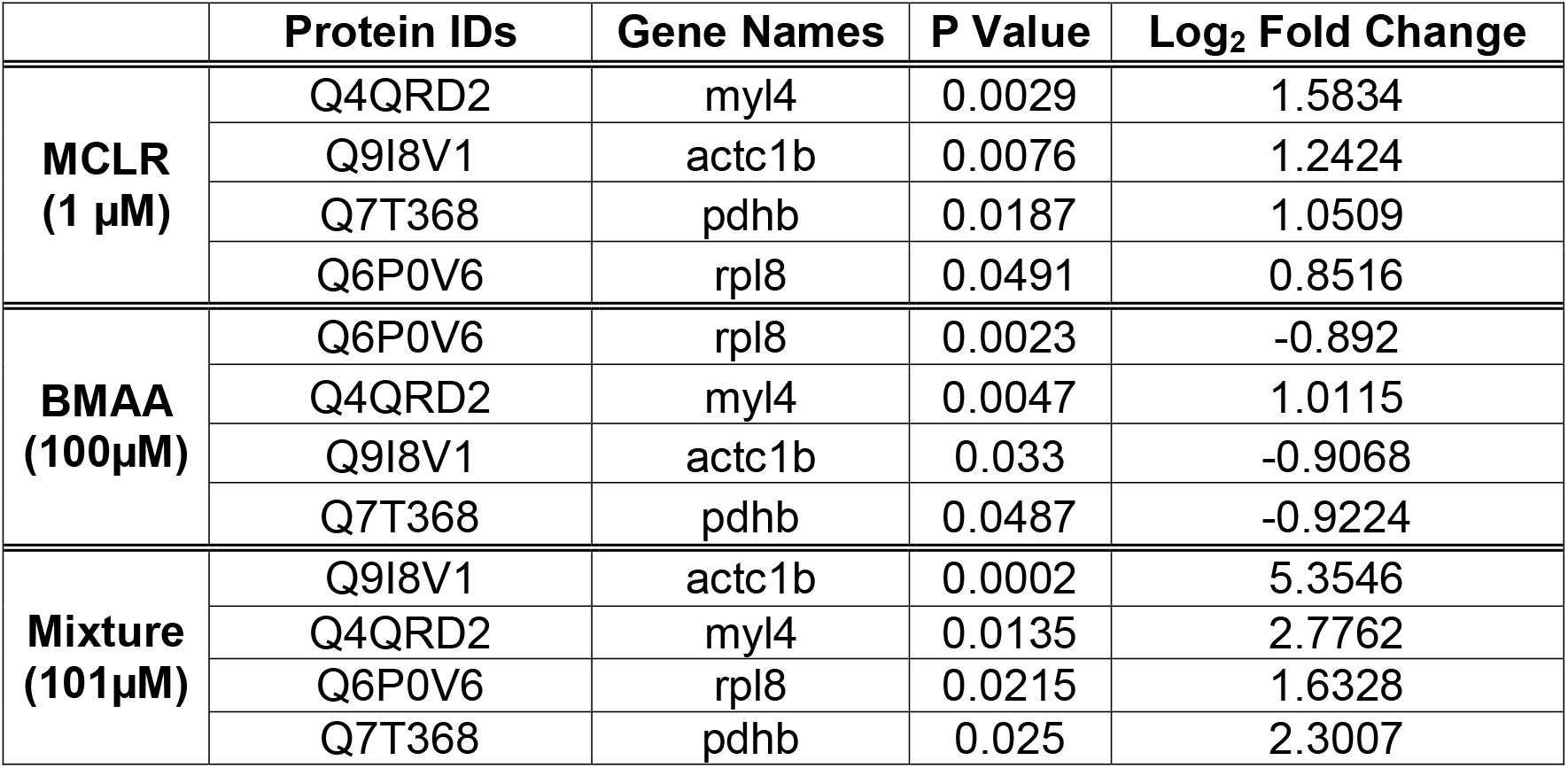
Shared DEPs amongst treatments.

|  | Protein IDs | Gene Names | P Value | Log<sub>2</sub> Fold Change |
| --- | --- | --- | --- | --- |
| **MCLR**<br>**(1 µM)** | Q4QRD2 | myl4 | 0.0029 | 1.5834 |
|  | Q9I8V1 | actc1b | 0.0076 | 1.2424 |
|  | Q7T368 | pdhb | 0.0187 | 1.0509 |
|  | Q6P0V6 | rpl8 | 0.0491 | 0.8516 |
| **BMAA**<br>**(100µM)** | Q6P0V6 | rpl8 | 0.0023 | -0.892 |
|  | Q4QRD2 | myl4 | 0.0047 | 1.0115 |
|  | Q9I8V1 | actc1b | 0.033 | -0.9068 |
|  | Q7T368 | pdhb | 0.0487 | -0.9224 |
| **Mixture**<br>**(101µM)** | Q9I8V1 | actc1b | 0.0002 | 5.3546 |
|  | Q4QRD2 | myl4 | 0.0135 | 2.7762 |
|  | Q6P0V6 | rpl8 | 0.0215 | 1.6328 |
|  | Q7T368 | pdhb | 0.025 | 2.3007 |

**Fig. 5.**
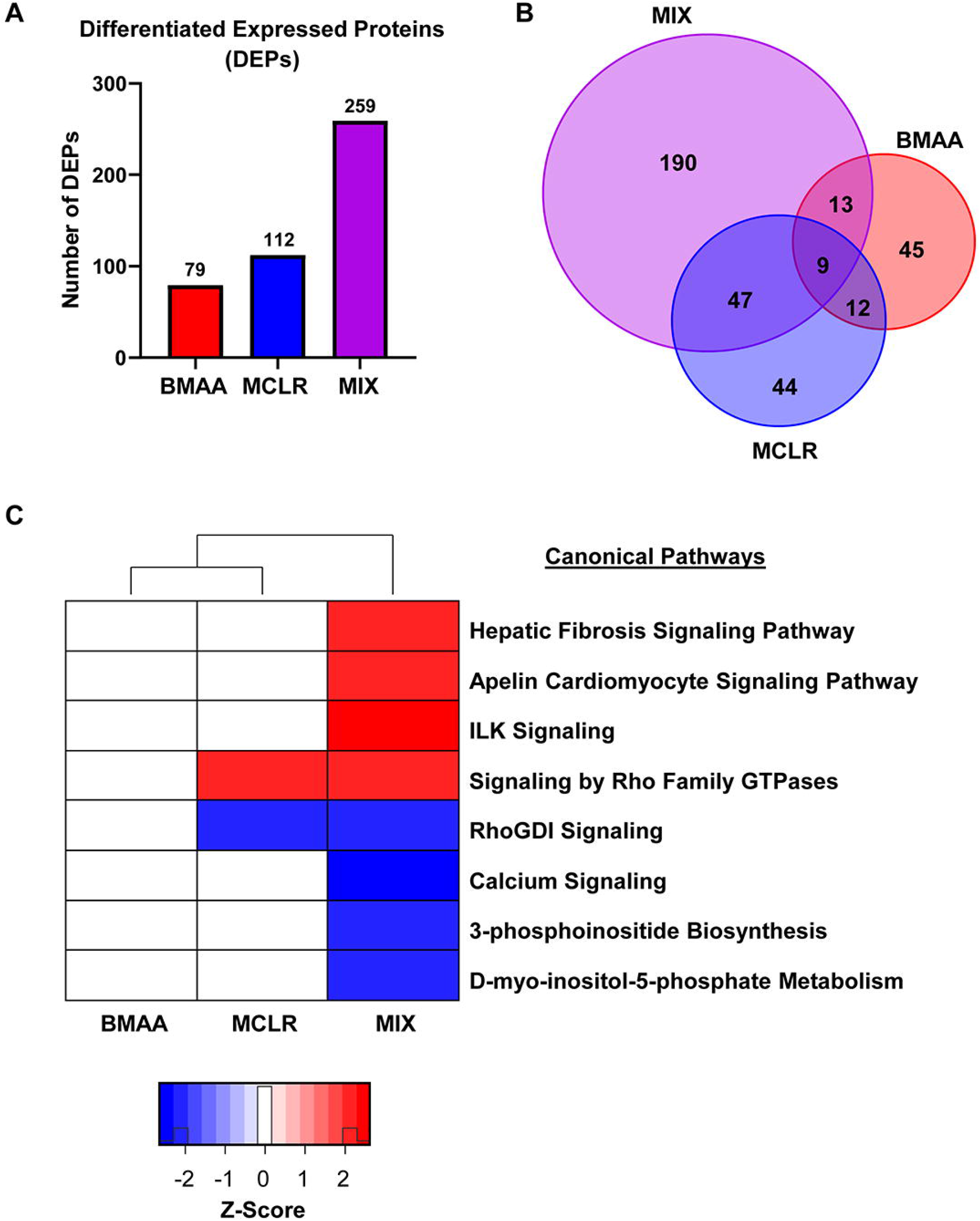
BMAA/MCLR mixture increases protein dysregulation *in vivo*. (A) Number of differentially expressed proteins (DEPs) per treatment condition. (B) Venn diagram showing overlap in DEPs between all treatment groups (red: BMAA protein group, the blue: MCLR protein group, purple: BMAA plus MCLR mixture protein group). (C) Heat map displaying the impacted canonical pathways from IPA functional analysis. The red or blue colored rectangles in each column indicates the z-score activities. Red shading indicates predicted activation and blue shading indicates predicted inhibition according to the scale at right. This heat map represents the z-scores obtained from the comparison between significantly expressed proteins (P≤◻0.05).

To further analyze the identified DEPs across treatments, we performed enrichment analysis to identify significantly perturbed pathways. **Supplemental table 5** lists all canonical pathways found to be significantly perturbed (z-score ≥|2|) along with their associated z-scores. Exposure to the BMAA/MCLR mixture enhanced the predicted activation/inhibition of eight canonical pathways (z-score≥|2|) compared to BMAA (zero) and MCLR (two), which supports the observation of a synergistic interaction *in vivo* (**Figure 5C**). RhoGDI signaling (z score=−2.121) and calcium signaling (z score=−2.449) were inhibited, while signaling by Rho Family GTPases (z-score=2.121) and ILK (z-score=2.121) were activated. Although exposure to MCLR at 1 μM did not cause behavioral modulation, two canonical pathways were significantly affected: 1) inhibition of RhoGDI signaling (z-score=−2), and 2) activation of signaling by Rho family GTPases (z-score=2) (**Figure 5C**). All four canonical pathways impacted by mixture exposure are broadly associated with neurotoxic processes related to reorganization of the actin cytoskeleton. Moreover, these pathway analysis results are reinforced by our protein interaction network analysis, in which we found significant differential regulation of key proteins associated with skeletal/muscular disorder and cellular assembly/organization (**Supplemental Table 6**). Within these networks, key neuronal proteins, including cell division cycle 42 (CDC-42; Enrichment p-value=0.0049; Log_2_ fold-change=1.8525), glutamate dehydrogenase 1 (GLUD1; Enrichment p-value=0.0299; Log_2_ fold-change= 2.6413), and the ALS-associated TDP-43 (TARDBP; Enrichment p-value=0.0299; Log_2_ fold-change= 2.5508) were significantly upregulated. Because TDP-43 is significantly associated with ALS disease pathology, we looked at expression of TDPBP and TDPBPL in all treatment groups. TDPBP was increased by MCLR exposure, but not BMAA exposure, and this increase was further enhanced by exposure to the BMAA/MCLR mixture (**Table 2**). Together, these data indicate that low concentrations of BMAA and MCLR in combination impact neurodegenerative processes in larval zebrafish.

**Table 2:** Impact of cyanotoxin exposure on TDP pathway proteins.

|  | Protein IDs | Gene Names | P Value | Log<sub>2</sub> Fold Change |
| --- | --- | --- | --- | --- |
| **BMAA**<br>**(100µM)** | Q802C7 | tardbp | 0.3614 | -1.3194 |
|  | Q6NYX2 | tardbpl | 0.9402 | 0.0674 |
| **MCLR**<br>**(1µM)** | Q802C7 | tardbp | 0.0245 | 1.6390 |
|  | Q6NYX2 | tardbpl | 0.7002 | 0.5127 |
| **Mixture**<br>**(101µM)** | Q802C7 | tardbp | 0.0159 | 2.5508 |
|  | Q6NYX2 | tardbpl | 0.0520 | 2.2426 |

## 4. Discussion

Since the 1950s, BMAA has been investigated for its potential to contribute to neurodegenerative diseases, including amyotrophic lateral sclerosis (ALS) (Reed et al. 1966). However, BMAA is only one of thousands of toxic metabolites produced by cyanobacteria (Dolman et al. 2012). While an increasing number of studies have singly addressed BMAA and its adverse effects, major knowledge gaps remain regarding the neuropathological effects of combined exposure to a cocktail of cyanotoxins. A number of studies have reported that cyanotoxins co-occur in natural environments (Banack et al. 2015; Jungblut et al. 2018; Metcalf et al. 2008; Sabart et al. 2015), including BMAA and the most abundant cyanotoxin, MCLR (McKindles et al. 2019). Although first considered to be primarily a hepatotoxin, MCLR has recently been shown to have neurotoxic effects both *in vitro* and *in vivo* (Li et al. 2012; Li et al. 2015; Wang et al. 2017; Wu et al. 2016). Thus, because BMAA and MCLR are ubiquitously present in the environment, have been previously detected together, and are neurotoxic, our study addresses the important question of whether they interact *in vivo* to enhance adverse effects. Building on prior work demonstrating that cyanotoxins can interact *in vitro* (Main et al. 2018; Martin et al. 2019), we show that 1) both BMAA and MCLR alter the behavior of larval zebrafish (**Figure 3**), 2) a simple binary mixture of BMAA and MCLR at low concentrations enhances behavioral neurotoxicity (**Figure 4**), and 3) BMAA and MCLR synergistically alter molecular changes associated with neuromuscular dysfunction (**Figure 5**).

Larval zebrafish have emerged as a powerful vertebrate model for studying neural development and behavioral circuits, as well as for translational toxicology (Tal et al. 2020; Wolman and Granato 2012). Although studying larvae is less directly relevant for studies of neurodegeneration, there is increasing evidence that developmental exposures can lead to disease later in life (Heindel and Vandenberg 2015). Indeed, neonatal exposure to BMAA has been found to cause motor defects and neurodegeneration in adult rats (Scott and Downing 2019; Scott et al. 2017). Thus, understanding the developmental impact of cyanotoxin exposure is critical for identifying potential early indicators of degenerative pathology. Here, we show that larval zebrafish behavior is modulated upon exposure to relatively low concentrations of both BMAA and MCLR in a dose-dependent manner. Although only traces of cyanotoxins have been found in large natural bodies of water (mean=41 μg.L^−1^) (Wiltsie et al. 2018), BMAA and MCLR can be found at relatively high concentrations (from ~0.02 to 8 mg.kg^−1^) in freshwater fish, crustaceans, and other types of seafood (Lance et al. 2018; Sahin et al. 1996) due to bioaccumulation through the food web. Thus, our mixture paradigm is an appropriate model of natural exposures.

Previous studies in larval zebrafish have indicated that BMAA may cause clonus like convulsions (Purdie et al. 2009) and pericardial edema and altered heart rate (Frøyset et al. 2016; Purdie et al. 2009). We did not observe these effects, but this could be due to differences in strain, embryo medium, exposure route, and analysis methods. In contrast to our data showing no effect on locomotion in bright light conditions (**Figure 2**), MCLR has previously been shown to reduce activity in zebrafish larvae in a light-dark assay (Tzima et al. 2017; Wu et al. 2016). This discrepancy could also arise from strain and media differences, but in agreement with these studies, we did not observe mortality or morphological defects in MCLR-exposed larvae. However, we detected significant, dose-dependent changes in acoustic startle behavior upon exposure to both BMAA and MCLR (**Figure 3**). These data reveal a need for greater standardization in zebrafish rearing methods, and they also show that our acoustic startle assay using high-speed cameras and kinematic analysis may be a broadly useful and highly sensitive addition to standard behavioral neurotoxicity testing.

The increased frequency of Mauthner-cell dependent short-latency startles (SLCs) in BMAA and MCLR-treated larvae indicates that the underlying sensorimotor circuit is hyperexcitable. BMAA is known to directly agonize glutamatergic receptors (Chiu et al. 2012; 2013), so the startle hypersensitivity in BMAA-treated fish could reflect that startle circuit neurons fire more easily following acoustic stimuli. Alternatively, hypersensitivity from exposure to these cyanotoxins could result from a reduction in inhibitory control of the startle circuit. Interestingly, both of these mechanisms have implications for ALS pathology, as excitotoxicity either from direct overstimulation of excitatory pathways, or from a loss of inhibitory input have been implicated in motor neuron death (Martin et al. 2012). Furthermore, our data show that BMAA and MCLR interact to enhance startle sensitivity at their respective NOAELs (**Figure 4**). To the best of our knowledge, only one previous study has examined the effects of exposure to BMAA and MCLR as a mixture. Anxiety-like behavior, exploratory behavior, and general locomotion were all found to be unchanged by acute exposure to a BMAA/MCLR mixture in the adult C57BL/6 mouse model (Myhre et al. 2018). This could indicate that the effects of the BMAA/MCLR mixture are limited to specific brain circuits, and/or that these neurotoxins exert their effects more strongly during early developmental stages (Karlsson et al. 2012; Scott et al. 2017). Future studies will examine the longer-term effects of developmental exposure to BMAA and MCLR.

To understand how BMAA and MCLR drive neurotoxicity, we used a label-free proteomics approach to identify the molecular pathways disrupted by BMAA/MCLR exposure in larval zebrafish. Our proteomics data display a clear trend towards enhanced toxicity in the mixture exposed group versus single exposures (**Figure 5A**), further supporting the conclusion from our behavioral data that the two toxins interact *in vivo*. Interestingly, DEPs displayed minimal overlap between treatments (**Figure 5B**), suggesting they act through different modes of action. It is notable that the BMAA/MCLR mixture impacted multiple critical cellular pathways, including signaling by ILK, Rho Family GTPases, RhoGDI, and calcium (**Figure 5C)**, which all impinge on regulation of the actin cytoskeleton. For example, overexpression of proteins in the Rho Family GTPase pathway such as CDC42 has specific effects on the actin filamentous system (Nobes and Hall 1995). CDC42 has a well-established role in triggering the formation/assembly of stress fibers mediated by Arp2/3-dependent actin nucleation (Aspenström 2019). These data are consistent with prior work showing that loss-of-function mutations in *cytoplasmic FMRP-interacting protein 2* (*cyfip2*), a key regulator of Arp2/3-mediated actin polymerization, cause startle hypersensitivity in larval zebrafish similar to that seen with BMAA/MCLR exposure (Marsden et al. 2018). In addition, previous reports show that MCLR induces neurotoxicity by triggering reorganization of actin cytoskeleton components (Li et al. 2012; Meng et al. 2011) by inhibiting serine/threonine-specific protein phosphatases (PPs) 1 and 2A (Huynh-Delerme et al. 2005; MacKintosh et al. 1990). Here, we show here that MCLR in combination with BMAA at low concentrations inhibits expression of these same protein phosphatases associated with cytoskeletal organization (PP1CAB (Q7ZVR3), enrichment p-value=0.0106; Log_2_ fold-change=−2.476; PP2CA (F1Q6Z7), enrichment p-value=0.0110; Log_2_ fold-change=−4.03; **Supplemental table 3**). Together, our molecular proteomics data support the idea that acoustic startle hypersensitivity may be an early indicator of neuronal stress.

That our unbiased proteomic analysis also revealed an upregulation of TDP-43 in BMAA/MCLR-exposed larvae (**Table 2, Supplemental Table 6**) is particularly striking. Cytoplasmatic TDP-43 inclusions are the key pathological hallmark in 98% of sALS cases (Mackenzie et al. 2010). Although our results cannot verify the sub-cellular localization of upregulated TDP-43, previous reports have shown that overexpression of TDP-43 in the cytoplasm leads to depletion of nuclear TDP-43, which has detrimental effects in mice (Fratta et al. 2018; Wils et al. 2010). While MCLR—but not BMAA— exposure also increased TDP-43 expression, this increase was exacerbated by the mixture, indicating that exposure to multiple cyanotoxins may enhance sALS disease processes. While the molecular mechanisms that specifically drive cyanotoxin-mediated neurotoxicity are not fully understood, our data support a model in which cyanotoxin mixtures cause neural dysfunction through multiple disease-associated pathways.

Together, our data provide new evidence that cyanotoxins synergistically interact *in vivo* to cause changes not only at the molecular level but also at the whole-organism level, as demonstrated by altered behavioral performance. Future work will seek to link specific molecular pathways, behavior regulation, and neuronal dysfunction, with the goal of revealing novel therapeutic and/or diagnostic targets for intractable neurodegenerative diseases such as ALS.

## Acknowledgments

We are thankful for startup funds provided by North Carolina State University (NCSU) and for pilot project support from the Center for Human Health and Environment (P30 ES025128). We also would like to thank Marsden and Bereman lab members for feedback on the manuscript. Finally, we are grateful to Derek Burton for zebrafish care and technical support for all experiments.

## Abbreviations

SLC: short latency c-startle
LLC: long latency c-startle
HPLC: high-performance liquid chromatography
LC/MS: high-pressure liquid chromatography combined mass spectrometry
IPA: ingenuity pathway analysis
ANOVA: analysis of variance
GO: gene ontology
BMAA: β-methylamino-L-alanine
MCLR: microcystin leucine-arginine
DEPs: differentially expressed proteins

## References

Aspenström P. 2019. The intrinsic gdp/gtp exchange activities of cdc42 and rac1 are critical determinants for their specific effects on mobilization of the actin filament system. Cells. 8(7).

Banack SA, Caller T, Henegan P, Haney J, Murby A, Metcalf JS, Powell J, Cox PA, Stommel E. 2015. Detection of cyanotoxins, β-n-methylamino-l-alanine and microcystins, from a lake surrounded by cases of amyotrophic lateral sclerosis. Toxins. 7(2):322–336.

Banack SA, Cox PA. 2003. Distribution of the neurotoxic nonprotein amino acid bmaa in cycas micronesica. Botanical Journal of the Linnean Society. 143(2):165–168.

Bereman MS, Beri J, Enders JR, Nash T. 2018. Machine learning reveals protein signatures in csf and plasma fluids of clinical value for als. Scientific reports. 8(1):16334–16314.

Beri J, Nash T, Martin RM, Bereman MS. 2017. Exposure to bmaa mirrors molecular processes linked to neurodegenerative disease. PROTEOMICS. 17(17-18):1700161-n/a.

Bozzoni V. 2016. Amyotrophic lateral sclerosis and environmental factors. Functional neurology. 31(1):7.

Bradley WG, Mash DC. 2009. Beyond guam: The cyanobacteria/bmaa hypothesis of the cause of als and other neurodegenerative diseases. Amyotrophic Lateral Sclerosis. 10(S2):7–20.

Brown RH, Al-Chalabi A. 2017. Amyotrophic lateral sclerosis. The New England Journal of Medicine. 377(2):162–172.

Burgess HA, Granato M. 2007. Modulation of locomotor activity in larval zebrafish during light adaptation. The Journal of experimental biology. 210(Pt 14):2526–2539.

Caller TA, Doolin JW, Haney JF, Murby AJ, West KG, Farrar HE, Ball A, Harris BT, Stommel EW. 2009. A cluster of amyotrophic lateral sclerosis in new hampshire: A possible role for toxic cyanobacteria blooms. Amyotrophic Lateral Sclerosis. 10(S2):101–108.

Chiu AS, Gehringer MM, Braidy N, Guillemin GJ, Welch JH, Neilan BA. 2012. Excitotoxic potential of the cyanotoxin β-methyl-amino-l-alanine (bmaa) in primary human neurons. Toxicon. 60(6):1159–1165.

Chiu AS, Gehringer MM, Braidy N, Guillemin GJ, Welch JH, Neilan BA. 2013. Gliotoxicity of the cyanotoxin, β-methyl-amino-l-alanine (bmaa). Scientific reports. 3(1):1482–1482.

Cox PA, Banack SA, Murch SJ, Rasmussen U, Tien G, Bidigare RR, Metcalf JS, Morrison LF, Codd GA, Bergman B et al. 2005. Diverse taxa of cyanobacteria produce β-n-methylamino-l-alanine, a neurotoxic amino acid. Proceedings of the National Academy of Sciences of the United States of America. 102(14):5074–5078.

Dolman AM, Rücker J, Pick FR, Fastner J, Rohrlack T, Mischke U, Wiedner C. 2012. Cyanobacteria and cyanotoxins: The influence of nitrogen versus phosphorus. PloS one. 7(6):e38757–e38757.

Field NC, Metcalf JS, Caller TA, Banack SA, Cox PA, Stommel EW. 2013. Linking β-methylamino-l-alanine exposure to sporadic amyotrophic lateral sclerosis in annapolis, md. Toxicon. 70:179–183.

Fratta P, Sivakumar P, Humphrey J, Lo K, Ricketts T, Oliveira H, Brito-Armas JM, Kalmar B, Ule A, Yu Y et al. 2018. Mice with endogenous tdp-43 mutations exhibit gain of splicing function and characteristics of amyotrophic lateral sclerosis. The EMBO Journal. 37(11):n/a-n/a.

Frøyset AK, Khan EA, Fladmark KE. 2016. Quantitative proteomics analysis of zebrafish exposed to sub-lethal dosages of β-methyl-amino-l-alanine (bmaa). Sci Rep. 6:29631.

Hao le T, Wolman M, Granato M, Beattie CE. 2012. Survival motor neuron affects plastin 3 protein levels leading to motor defects. J Neurosci. 32(15):5074–5084.

Heindel JJ, Vandenberg LN. 2015. Developmental origins of health and disease: A paradigm for understanding disease cause and prevention. Curr Opin Pediatr. 27(2):248–253.

Huynh-Delerme C, Edery M, Huet H, Puiseux-Dao S, Bernard C, Fontaine J-J, Crespeau F, de Luze A. 2005. Microcystin-lr and embryo–larval development of medaka fish, oryzias latipes. I. Effects on the digestive tract and associated systems. Toxicon. 46(1):16–23.

Ingre C, Roos PM, Piehl F, Kamel F, Fang F. 2015. Risk factors for amyotrophic lateral sclerosis. Clinical epidemiology. 7:181–193.

Jones N. 2009. Genetic risk factors for sporadic als. Nature Reviews Neurology. 5(11):579–579.

Jungblut AD, Wilbraham J, Banack SA, Metcalf JS, Codd GA. 2018. Microcystins, bmaa and bmaa isomers in 100-year-old antarctic cyanobacterial mats collected during captain r.F. Scott’s discovery expedition. European Journal of Phycology. 53(2):115.

Karlsson O, Berg A-L, Lindström A-K, Hanrieder J, Arnerup G, Roman E, Bergquist J, Lindquist NG, Brittebo EB, Andersson M et al. 2012. Neonatal exposure to the cyanobacterial toxin bmaa induces changes in protein expression and neurodegeneration in adult hippocampus. Toxicological sciences : an official journal of the Society of Toxicology. 130(2):391–404.

Karlsson O, Michno W, Ransome Y, Hanrieder J. 2017. Maldi imaging delineates hippocampal glycosphingolipid changes associated with neurotoxin induced proteopathy following neonatal bmaa exposure. BBA – Proteins and Proteomics. 1865(7):740–746.

Khatri P, Drăghici S. 2005. Ontological analysis of gene expression data: Current tools, limitations, and open problems. Bioinformatics (Oxford, England). 21(18):3587–3595.

Kurland LT, Mulder DW. 1955. Epidemiologic investigations of amyotrophic lateral sclerosis. 2. xsFamilial aggregations indicative of dominant inheritance. Ii. Neurology. 5(4):249.

Lance E, Arnich N, Maignien T, Biré R. 2018. Occurrence of β-n-methylamino-l-alanine (bmaa) and isomers in aquatic environments and aquatic food sources for humans. Switzerland: MDPI AG. p. 83.

Li G, Cai F, Yan W, Li C, Wang J. 2012. A proteomic analysis of mclr-induced neurotoxicity: Implications for alzheimer’s disease. Toxicological sciences : an official journal of the Society of Toxicology. 127(2):485–495.

Li G, Yan W, Dang Y, Li J, Liu C, Wang J. 2015. The role of calcineurin signaling in microcystin-lr triggered neuronal toxicity. Scientific reports. 5(1):11271–11271.

Lobner D, Piana PMT, Salous AK, Peoples RW. 2006. Β-n-methylamino-l-alanine enhances neurotoxicity through multiple mechanisms. Neurobiology of Disease. 25(2):360–366.

Mackenzie IR, Rademakers R, Neumann M. 2010. Tdp-43 and fus in amyotrophic lateral sclerosis and frontotemporal dementia. Lancet Neurol. 9(10):995–1007.

MacKintosh C, Beattie KA, Klumpp S, Cohen P, Codd GA. 1990. Cyanobacterial microcystin-lr is a potent and specific inhibitor of protein phosphatases 1 and 2a from both mammals and higher plants. FEBS Letters. 264(2):187–192.

Main BJ, Main BJ, Rodgers KJ, Rodgers KJ. 2018. Assessing the combined toxicity of bmaa and its isomers 2,4-dab and aeg in vitro using human neuroblastoma cells. Neurotoxicity Research. 33(1):33–42.

Marquart GD, Tabor KM, Bergeron SA, Briggman KL, Burgess HA. 2019. Prepontine non-giant neurons drive flexible escape behavior in zebrafish. PLOS Biology. 17(10):e3000480–e3000480.

Marsden KC, Jain RA, Wolman MA, Echeverry FA, Nelson JC, Hayer KE, Miltenberg B, Pereda AE, Granato M. 2018. A cyfip2-dependent excitatory interneuron pathway establishes the innate startle threshold. Cell reports. 23(3):878–887.

Martin LJ, Martin LJ, Chang Q, Chang Q. 2012. Inhibitory synaptic regulation of motoneurons: A new target of disease mechanisms in amyotrophic lateral sclerosis. Molecular Neurobiology. 45(1):30–42.

Martin RM, Stallrich J, Bereman MS. 2019. Mixture designs to investigate adverse effects upon co-exposure to environmental cyanotoxins. Toxicology. 421:74–83.

Masseret E, Banack S, Boumédiène F, Abadie E, Brient L, Pernet F, Juntas-Morales R, Pageot N, Metcalf J, Cox P et al. 2013. Dietary bmaa exposure in an amyotrophic lateral sclerosis cluster from southern france. PloS one. 8(12):e83406–e83406.

McKindles KM, Zimba PV, Chiu AS, Watson SB, Gutierrez DB, Westrick J, Kling H, Davis TW. 2019. A multiplex analysis of potentially toxic cyanobacteria in lake winnipeg during the 2013 bloom season. Toxins. 11(10):587.

Meng G, Sun Y, Fu W, Guo Z, Xu L. 2011. Microcystin-lr induces cytoskeleton system reorganization through hyperphosphorylation of tau and hsp27 via pp2a inhibition and subsequent activation of the p38 mapk signaling pathway in neuroendocrine (pc12) cells. Ireland: Elsevier Ireland Ltd. p. 218–229.

Metcalf JS, Banack SA, Lindsay J, Morrison LF, Cox PA, Codd GA. 2008. Co-occurrence of β-n-methylamino-l-alanine, a neurotoxic amino acid with other cyanobacterial toxins in british waterbodies, 1990–2004. Oxford, UK: Blackwell Publishing Ltd. p. 702–708.

Metcalf JS, Richer R, Cox PA, Codd GA. 2012. Cyanotoxins in desert environments may present a risk to human health. Science of the Total Environment. 421-422:118–123.

Myhre O, Eide DM, Kleiven S, Utkilen HC, Hofer T. 2018. Repeated five-day administration of l-bmaa, microcystin-lr, or as mixture, in adult c57bl/6 mice – lack of adverse cognitive effects. Scientific reports. 8(1):2308–2314.

Nobes CD, Hall A. 1995. Rho, rac, and cdc42 gtpases regulate the assembly of multimolecular focal complexes associated with actin stress fibers, lamellipodia, and filopodia. Cell. 81(1):53–62.

Purdie EL, Samsudin S, Eddy FB, Codd GA. 2009. Effects of the cyanobacterial neurotoxin β-n-methylamino-l-alanine on the early-life stage development of zebrafish (danio rerio). Aquatic Toxicology. 95(4):279–284.

Reed D, Plato C, Elizan T, Kurland LT. 1966. The amyotrophic lateral sclerosis/parkinsonism-dementia complex: A ten-year follow-up on guam. I. Epidemiologic studies. United States. p. 54.

Renton AE, Chiò A, Traynor BJ. 2014. State of play in amyotrophic lateral sclerosis genetics. Nature neuroscience. 17(1):17–23.

Sabart M, Crenn K, Perrière F, Abila A, Leremboure M, Colombet J, Jousse C, Latour D. 2015. Co-occurrence of microcystin and anatoxin-a in the freshwater lake aydat (france): Analytical and molecular approaches during a three-year survey. Harmful Algae. 48:12–20.

Sahin A, Tencalla FG, Dietrich DR, Naegeli H. 1996. Biliary excretion of biochemically active cyanobacteria (blue-green algae) hepatotoxins in fish. Ireland: Elsevier Ireland Ltd. p. 123–130.

Scott L, Downing T. 2019. Dose-dependent adult neurodegeneration in a rat model after neonatal exposure to β-n-methylamino-l-alanine. Neurotoxicity Research. 35(3):711–723.

Scott LL, Downing TG, Timothy D, Laura S. 2017. A single neonatal exposure to bmaa in a rat model produces neuropathology consistent with neurodegenerative diseases. Toxins. 10(1):22.

Swinnen B, Robberecht W. 2014. The phenotypic variability of amyotrophic lateral sclerosis. Nature reviews Neurology. 10(11):661–670.

Tal T, Yaghoobi B, Lein PJ. 2020. Translational toxicology in zebrafish. Current opinion in toxicology.

Tyanova S, Temu T, Sinitcyn P, Carlson A, Hein MY, Geiger T, Mann M, Cox J. 2016. The perseus computational platform for comprehensive analysis of (prote)omics data. Nature methods. 13(9):731–740.

Tzima E, Serifi I, Tsikari I, Alzualde A, Leonardos I, Papamarcaki T. 2017. Transcriptional and behavioral responses of zebrafish larvae to microcystin-lr exposure. International journal of molecular sciences. 18(2):365.

Wang B, Liu J, Huang P, Xu K, Wang H, Wang X, Guo Z, Xu L. 2017. Protein phosphatase 2a inhibition and subsequent cytoskeleton reorganization contributes to cell migration caused by microcystin-lr in human laryngeal epithelial cells (hep-2). United States: Wiley Subscription Services, Inc. p. 890–903.

Wang Q, Xie P, Chen J, Liang G. 2008. Distribution of microcystins in various organs (heart, liver, intestine, gonad, brain, kidney and lung) of wistar rat via intravenous injection. Toxicon. 52(6):721–727.

Wils H, Kleinberger G, Janssens J, Pereson S, Joris G, Cuijt I, Smits V, Ceuterick-de Groote C, Van Broeckhoven C, Kumar-Singh S. 2010. Tdp-43 transgenic mice develop spastic paralysis and neuronal inclusions characteristic of als and frontotemporal lobar degeneration. Proc Natl Acad Sci U S A. 107(8):3858–3863.

Wiltsie D, Schnetzer A, Green J, Vander Borgh M, Fensin E. 2018. Algal blooms and cyanotoxins in jordan lake, north carolina. Toxins. 10(2):92.

Wolman M, Granato M. 2012. Behavioral genetics in larval zebrafish: Learning from the young. Hoboken: Wiley Subscription Services, Inc., A Wiley Company. p. 366–372.

Wu Q, Yan W, Liu C, Li L, Yu L, Zhao S, Li G. 2016. Microcystin-lr exposure induces developmental neurotoxicity in zebrafish embryo. Environmental Pollution. 213:793–800.

Zhao S, Li G, Chen J. 2015. A proteomic analysis of prenatal transfer of microcystin-lr induced neurotoxicity in rat offspring. J Proteomics. 114:197–213.

