## Supplemental Information for "Exposure to a mixture of BMAA and MCLR synergistically modulates behavior in larval zebrafish while exacerbating molecular changes related to neurodegeneration"

Submitted to: Toxicological Sciences

Submitted:

Pages: 13

Supplemental Figures: 2

Supplemental Tables: 6

Running Title: Investigation of interactions amongst BMAA and MCLR *in vivo*.

Keywords: Cyanotoxins; Mixtures; Synergism; Zebrafish; Behavior; Proteomics

*Author for Correspondence

Kurt C. Marsden

Department of Biological Sciences

North Carolina State University

Raleigh, NC

**Supplemental Figure Legends**

**Supplemental Figure 1. Determination of routine swims and turns.** (A-B) Violin plots depict the frequency distribution of spontaneous turns and swims, respectively. Levels not connected by the same letter are significantly different, Tukey-Kramer HSD, Alpha 0.05).

**Supplemental Figure 2**. **Volcano plots.** (A-C) Volcano plots representation of BMAA (100µM), MCLR(1µM), and BMAA/MCLR mixture (101µM) regulated proteins. Red, blue, and purple dots represent significantly (p ≤ 0.05) regulated proteins (BMAA, MCLR, and BMAA/MCLR mixture, respectively) while black dot represents non-significantly (p ≥ 0.05) proteins compared to control.

**Supplemental Methods**

**Sample Collection, Protein Extraction and Digestion**

Sample Collection – Three biological replicate were prepared for each treat ment condition where samples were carefully collected and flash frozen using liquid nitrogen and stored at -80 °F.

Protein Extraction - A pool of 10 zebrafish larvae whole body for each condition were suspended in a 100 µL solution of 50 mM ammonium bicarbonate (pH 8.0) containing 1% SDC. Zebrafish bodies were lysed by probe sonication via 2 pulses at 20 seconds per pulse at a 20% amplitude setting. Cell debris was removed via centrifugation at 10,000 RPM for 5 minutes. The supernatant was retained and assessed for protein quantification by bicinchoninic acid (BCA) assay.

Protein digestion – 100 ug of protein was diluted to a final volume of 100 µL with the 50 mM ammonium bicarbonate and 1% SDC solution. DTT was added to a final concentration of 5 mM and incubated at 60ºC for 30 minutes to reduce disulfide bonds. Samples were cooled to room temperature, followed by alkylation of cysteine residues with iodoacetamide (15 mM) at room temperature and in the dark for 20 minutes. Samples were then subjected to filter-aided sample preparation (FASP) (Wisniewski et al. 2009) for protein cleanup using Vivacon 30,000 kDa molecular weight cutoff filters and reconstituted with 100 µL of 50 mM ammonium bicarbonate solution. Tryptic digestion was carried out by hydrating lyophilized trypsin to a stock solution of 1 µg/µL with 0.01% acetic acid in water followed by addition of trypsin to the protein mixture at a 1:50 ratio and then incubated at 37ºC for 4 hours. After digestion, peptides were acidified with HCl at a final concentration of 250 mM (pH ≤ 3).

**LC-MS/MS Analysis**

3 µL of each sample were injected and analyzed using an Easy nanoLC 1200 coupled to an Orbitrap Exploris Mass Spectrometer (Thermo Scientific, Bremen, Germany). LC separation was performed with an EASY-spray system, consisting of a 50 cm, 75 μm ID PepMap RSLC, C18, 100 Å, 2μm particles, in which was connected to an Easy-nLC Ultra UHPLC system (Thermo Scientific, Odense, DK). Separation of peptides was achieved through a gradient of mobile phase A (98% water, 2% acetonitrile, and 0.1% formic acid) and mobile phase B (99.9% acetonitrile and 0.1% formic acid). The LC method applied consisted of a gradient of 5%-50% B over 120 minutes, followed by a ramp to 95% B in two minute. The column was washed at 95% B for 16 minutes. Tandem mass spectrometry was carried out using positive ion mode and data dependent mode via cycling every 3 seconds (i.e., master scan time). MS1 scans were performed at a resolving power of 60,000 from m/z 375 to 1600 and an automatic gain control (AGC) target of 1 x 10^6. MS2 scans were performed at a resolving power of 15,000 and an AGC target of 5 x 10^4. A 30 second dynamic exclusion window was applied during sampling to avoid repeated interrogation of high-abundant species.

Wisniewski, J.R., Zougman, A., Nagaraj, N. and Mann, M. 2009. Universal sample preparation method for proteome analysis. Nat Methods 6, 359-U360.

|  | **Protein IDs** | **Gene names** | **P Value** | **Fold Change** |
| --- | --- | --- | --- | --- |
| **Mix** | Q9I8V1 | actc1b | 0.0002 | 5.3546 |
|  | G1K2X0 | ttnb | 0.0075 | 0.4399 |
|  | F1Q4Q4 | lmf2b | 0.0114 | 2.0293 |
|  | Q4QRD2 | myl4 | 0.0135 | 2.7762 |
|  | Q6P0V6 | rpl8 | 0.0215 | 1.6328 |
|  | Q7T368 | pdhb | 0.0250 | 2.3007 |
|  | A2BGE0 | stxbp1b | 0.0460 | -1.1152 |
| **MCLR** | Q4QRD2 | myl4 | 0.0029 | 1.5834 |
|  | F1Q4Q4 | lmf2b | 0.0075 | 1.4327 |
|  | Q9I8V1 | actc1b | 0.0076 | 1.2424 |
|  | Q7T368 | pdhb | 0.0187 | 1.0509 |
|  | G1K2X0 | ttnb | 0.0306 | 0.2465 |
|  | A2BGE0 | stxbp1b | 0.0463 | -0.8948 |
|  | Q6P0V6 | rpl8 | 0.0491 | 0.8516 |
| **BMAA** | Q6P0V6 | rpl8 | 0.0023 | -0.8920 |
|  | Q4QRD2 | myl4 | 0.0047 | 1.0115 |
|  | F1Q4Q4 | lmf2b | 0.0157 | 1.6026 |
|  | A2BGE0 | stxbp1b | 0.0280 | -0.4347 |
|  | Q9I8V1 | actc1b | 0.0330 | -0.9068 |
|  | G1K2X0 | ttnb | 0.0386 | -0.3416 |
|  | Q7T368 | pdhb | 0.0487 | -0.9224 |

**Supplemental Table 1: DEPs shared in all treatments.** This table lists the gene name, p value and fold change for all seven DEPs under each treatment group: mixture, MCLR , and BMAA (cut-off p value of 0.05).

| **Protein ID** | **P value** | **Fold change** |
| --- | --- | --- |
| P61485 | 0.000537997 | 1.03133 |
| F6NPB1 | 0.001029248 | -0.724581 |
| Q6PGZ2 | 0.001578629 | -1.38599 |
| Q6PH70 | 0.001604132 | -0.697016 |
| Q7ZUI0 | 0.002883899 | 1.71248 |
| Q4QRD2 | 0.002940424 | 1.58339 |
| Q6PBH7 | 0.003666655 | 0.920369 |
| Q1LWN2 | 0.005450169 | 7.87221 |
| Q7ZUB3 | 0.005715576 | -0.697507 |
| Q498X8 | 0.005717419 | 0.294402 |
| Q5XJX0 | 0.006209691 | -2.29377 |
| F1QLA4 | 0.006449262 | 1.74926 |
| Q1LVF8 | 0.007009709 | -1.34431 |
| F1QKE8 | 0.007318634 | 1.70647 |
| Q7SY49 | 0.007508791 | 0.654617 |
| F1Q4Q4 | 0.007509828 | 1.43271 |
| Q9I8V1 | 0.007594514 | 1.24242 |
| X1WC29 | 0.007737488 | 2.07292 |
| Q568G0 | 0.007747293 | 4.3661 |
| Q6DRJ2 | 0.008702018 | -0.522572 |
| E7FBV0 | 0.009285601 | -1.7721 |
| F1Q7H4 | 0.010566958 | -0.333159 |
| C5NU62 | 0.010836275 | 0.604985 |
| E7F1C8 | 0.011581638 | -0.588008 |
| Q1MTC4 | 0.011717636 | 5.64942 |
| Q499A8 | 0.012082311 | 0.877755 |
| F1RD80 | 0.012321115 | 1.53995 |
| Q6NY49 | 0.01242682 | 0.296485 |
| A8DZ95 | 0.012677394 | 0.739624 |
| E9QF69 | 0.013574692 | 0.700799 |
| H0WEA8 | 0.013649288 | -3.05229 |
| B0UY94 | 0.013687685 | 1.97797 |
| F1QPE9 | 0.014507414 | -1.6243 |
| Q7ZUD3 | 0.014974764 | 0.978234 |
| F1RCL9 | 0.015219837 | -1.52614 |
| Q66HY3 | 0.01555894 | -0.157012 |
| B8A568 | 0.015637236 | 0.323329 |
| B0S6K5 | 0.017210362 | -0.42228 |
| Q1L8L9 | 0.017232171 | 1.44158 |
| Q6WFZ4 | 0.017670547 | 0.229356 |
| Q6PFT4 | 0.017911007 | -0.609681 |
| Q5U396 | 0.017968425 | 0.37271 |
| Q804H2 | 0.018337072 | 2.69923 |
| Q5RHV1 | 0.018337495 | -0.530611 |
| Q7T368 | 0.018726215 | 1.05092 |
| Q7ZYX4 | 0.018781056 | 1.59713 |
| B8JMZ1 | 0.019126258 | 2.25584 |
| Q7T2C5 | 0.019268149 | -1.12868 |
| Q6DBX3 | 0.019822572 | 0.510787 |
| K7DYE9 | 0.019981588 | -2.54357 |
| A8KB78 | 0.020442782 | 1.28981 |
| H0WF22 | 0.0218454 | 2.06132 |
| B5DE37 | 0.022747308 | 0.286835 |
| E9QG51 | 0.023196369 | 1.09185 |
| Q802C7 | 0.024514893 | 1.63895 |
| Q8AYH1 | 0.026836755 | 0.959012 |
| Q2YDS4 | 0.0278157 | -0.502092 |
| Q52PJ7 | 0.028102179 | 1.64908 |
| F1QX79 | 0.028536454 | -1.952 |
| Q802F5 | 0.02898211 | -0.593355 |
| B0JZP9 | 0.029125943 | -0.91486 |
| Q802Z4 | 0.029201825 | -2.22581 |
| R4GEM8 | 0.02971666 | 0.22159 |
| E7F605 | 0.029787908 | 0.481248 |
| G1K2X0 | 0.030550618 | 0.246496 |
| Q568E8 | 0.030846081 | -0.412317 |
| B2GSC6 | 0.033874274 | 1.79971 |
| F1R657 | 0.034515169 | -0.578229 |
| Q6P0G2 | 0.034542996 | 0.987933 |
| F1RDE6 | 0.034870647 | -0.700619 |
| F1RBG8 | 0.035580332 | -0.559079 |
| F1QXG4 | 0.036166793 | 2.54866 |
| Q7ZVY9 | 0.036466157 | 1.35228 |
| B2CZC4 | 0.03672823 | -0.278074 |
| E9QFD8 | 0.037401584 | 2.18335 |
| Q7SXT4 | 0.037732015 | 1.10818 |
| X1WEV8 | 0.038104828 | 0.472936 |
| Q4KMD5 | 0.038433506 | -0.748683 |
| Q08CE4 | 0.038596685 | -1.35093 |
| F1QS02 | 0.03908499 | -1.32183 |
| I3IRW5 | 0.040209623 | 0.55971 |
| F1R1C0 | 0.040488343 | 1.30374 |
| R4GED4 | 0.040854509 | -0.814199 |
| Q6NXC1 | 0.041061991 | 1.73685 |
| Q502B2 | 0.042491298 | -0.75292 |
| X1WFW6 | 0.042825259 | -1.74899 |
| Q7T2P7 | 0.04336806 | 2.65093 |
| Q6PFU7 | 0.043597341 | 1.11463 |
| E7F4R0 | 0.043952137 | -1.06001 |
| F1QSR4 | 0.044132649 | -0.634272 |
| Q0V948 | 0.044137731 | 1.19904 |
| F1QKF9 | 0.044180436 | 0.850011 |
| Q7ZZW4 | 0.044522547 | 0.604935 |
| E7F0A1 | 0.045000776 | 1.21869 |
| F1QEW4 | 0.045823684 | 0.717285 |
| Q7SX99 | 0.046131757 | 2.74944 |
| A2BGE0 | 0.046288169 | -0.894755 |
| Q1MTC6 | 0.046311623 | 2.68835 |
| F1Q909 | 0.046314822 | 1.23109 |
| B0UYB4 | 0.046895374 | -1.11569 |
| E7EY75 | 0.047272656 | 1.49723 |
| A9C3T0 | 0.047384903 | 0.863347 |
| Q0P3Z4 | 0.047789226 | 0.700256 |
| Q6TNV6 | 0.048071757 | 1.60884 |
| Q66I22 | 0.048400517 | -0.896641 |
| Q5RGI5 | 0.049086266 | -0.624488 |
| Q6DRK0 | 0.049098701 | -0.407731 |
| Q6P0V6 | 0.049147338 | 0.85161 |
| Q567N9 | 0.049151865 | -0.567569 |
| E7F134 | 0.049688971 | 0.861015 |
| Q5U3U8 | 0.04989649 | 0.355696 |
| F8W2B6 | 0.049933269 | 2.17735 |

**Supplemental Table 2:** Differentiated expressed proteins for the MCLR (1µM) exposed group.

| **Protein IDs** | **P value** | **Fold Change** |
| --- | --- | --- |
| E7F689 | 8.4561E-05 | -1.53001 |
| F1R657 | 0.000117961 | -1.47213 |
| Q9I8V1 | 0.0002142 | 5.35456 |
| E9QG51 | 0.000440677 | 1.91464 |
| Q803J5 | 0.000817881 | -1.92007 |
| F1QCU4 | 0.001248418 | -1.82011 |
| B8A568 | 0.001290298 | 1.0146 |
| Q0IIQ6 | 0.001769661 | -1.96776 |
| Q4V9E0 | 0.002005857 | 2.61601 |
| Q7ZYX4 | 0.002326002 | 3.68173 |
| E7FBV0 | 0.002503341 | -1.75537 |
| B0R0B3 | 0.002636999 | -1.52612 |
| B8JMZ1 | 0.002863782 | 3.0247 |
| F1RD80 | 0.003288365 | 3.32832 |
| Q4KMJ5 | 0.003884007 | -2.16949 |
| E9QF69 | 0.004055459 | 1.50802 |
| Q6DHC7 | 0.00420746 | -1.01721 |
| Q7SZR6 | 0.004333712 | 1.38333 |
| B2GSC6 | 0.004437823 | 4.11901 |
| Q0P3Z4 | 0.004546425 | 1.4931 |
| A8WG05 | 0.004787403 | 1.82707 |
| B0UXI4 | 0.004944814 | 2.1441 |
| Q52WY2 | 0.004992522 | 1.85253 |
| Q5W7F1 | 0.005207509 | -0.813051 |
| Q6PBH7 | 0.005512137 | 2.26218 |
| A8KAZ5 | 0.005683947 | -2.23491 |
| P61485 | 0.005810587 | 1.81606 |
| Q568G0 | 0.005863001 | 5.02741 |
| Q7ZUG5 | 0.006017554 | 2.0337 |
| B8JKT0 | 0.006043799 | 2.19509 |
| F1Q9K4 | 0.00608023 | -0.609554 |
| H9GXK5 | 0.00611998 | -2.11196 |
| F1QX55 | 0.006776259 | -2.11314 |
| E7EY75 | 0.007137235 | 3.50116 |
| Q5XJD1 | 0.007252538 | -0.376392 |
| E7FGL2 | 0.007318296 | -3.05072 |
| Q804H2 | 0.007339391 | 4.10424 |
| G1K2X0 | 0.00748652 | 0.43993 |
| Q52PJ7 | 0.007740696 | 3.27734 |
| F1QIP3 | 0.008071793 | -1.65917 |
| Q6P960 | 0.0081313 | -2.21665 |
| F1Q909 | 0.008136357 | 3.45387 |
| Q6NWE6 | 0.00854555 | -2.46912 |
| F1QTQ2 | 0.008606964 | -1.24617 |
| Q4VBV1 | 0.008766576 | 0.853519 |
| F1R886 | 0.009098923 | -2.13874 |
| F1QD76 | 0.0091262 | -1.65043 |
| Q802Z4 | 0.009175506 | -1.38899 |
| X1WC29 | 0.009539596 | 2.13454 |
| B0S5K4 | 0.009964603 | 2.61194 |
| F1QZL8 | 0.010171617 | -1.80227 |
| A1L1F7 | 0.010190606 | -1.09439 |
| Q502C8 | 0.010392739 | 3.41457 |
| F1QRV5 | 0.010466705 | -2.54144 |
| Q0PZF1 | 0.010628207 | 3.88837 |
| Q7ZVR3 | 0.010649029 | -2.47598 |
| B0UY94 | 0.010884789 | 3.75808 |
| F1Q6Z7 | 0.011023766 | -4.03002 |
| Q06W26 | 0.011340965 | -1.78242 |
| F1Q4Q4 | 0.01144353 | 2.02926 |
| F1RC03 | 0.011449855 | 0.45544 |
| Q1LUC1 | 0.011873534 | -1.46352 |
| A1L1W9 | 0.01196961 | -1.08241 |
| Q6P0G2 | 0.012000795 | 1.78233 |
| E7F8C0 | 0.012468666 | -0.657375 |
| Q6PGZ2 | 0.013040072 | -2.31363 |
| Q4QRD2 | 0.013451168 | 2.77618 |
| H9GZ72 | 0.013451478 | -1.97242 |
| Q6DI16 | 0.013654946 | 2.43357 |
| Q6DRJ2 | 0.013783198 | -0.729303 |
| Q6NXC1 | 0.013825791 | 3.52501 |
| Q7ZTZ7 | 0.014087697 | -0.568653 |
| Q8AW03 | 0.014109122 | 1.94919 |
| F1Q7H4 | 0.01414979 | -0.558552 |
| Q1L8L9 | 0.014237705 | 2.12686 |
| P83571 | 0.014469383 | 3.008 |
| F1Q615 | 0.014810851 | 4.28158 |
| Q6DGN1 | 0.015047003 | 2.43111 |
| E7FC33 | 0.015106714 | -1.89419 |
| F1QKE8 | 0.015511364 | 1.98637 |
| F1Q8K1 | 0.015821585 | 1.2054 |
| Q7ZZW4 | 0.015859153 | 1.23798 |
| F1QBK3 | 0.015897177 | -1.42172 |
| Q802C7 | 0.015899007 | 2.55077 |
| Q6DC41 | 0.015930155 | 0.654884 |
| Q7SYL6 | 0.016207275 | -2.19961 |
| Q7ZUV8 | 0.016284709 | -2.39615 |
| F1QRG5 | 0.016409298 | -2.41936 |
| Q7T2P7 | 0.01650517 | 3.71871 |
| Q567N1 | 0.017111576 | 2.28231 |
| Q7ZUD3 | 0.017423685 | 0.917695 |
| F6NPB1 | 0.017687237 | -0.977498 |
| Q9W6T5 | 0.017699459 | 0.963429 |
| B3DJ34 | 0.017897402 | 0.558185 |
| F1QS02 | 0.018042637 | -1.99283 |
| A9JRY8 | 0.018206229 | 1.03136 |
| A8E7G5 | 0.018347631 | 0.975543 |
| Q803D2 | 0.018757286 | 0.914983 |
| Q6NYP0 | 0.018758582 | -1.622 |
| F1R1T1 | 0.018929973 | -0.943645 |
| X1WHD9 | 0.019042765 | 2.53941 |
| A8E5C5 | 0.019137711 | 2.28742 |
| Q5PR98 | 0.01944957 | 4.02381 |
| Q803B0 | 0.019529 | 3.58377 |
| E7F2J3 | 0.019550146 | 1.50016 |
| F1QP57 | 0.019582133 | -1.36679 |
| H0WF22 | 0.019590702 | 2.33149 |
| Q499A8 | 0.019766055 | 2.17604 |
| B0S6K5 | 0.019995396 | -0.737628 |
| X1WCJ5 | 0.020113609 | -2.11764 |
| A4QP54 | 0.020189706 | 2.10885 |
| F1R4G7 | 0.020212033 | 1.32093 |
| Q568R7 | 0.020458322 | -0.943204 |
| Q6P972 | 0.020681879 | 2.66752 |
| Q6TNS5 | 0.020793136 | 0.76698 |
| A8WFU6 | 0.020863636 | 2.57714 |
| Q6PCS7 | 0.020980218 | -3.24506 |
| Q5RHR9 | 0.021086767 | 1.87021 |
| Q6P0V6 | 0.021536246 | 1.63275 |
| T1ECV3 | 0.021575954 | 2.58773 |
| Q6IQH2 | 0.021627683 | -2.68027 |
| A8BBH7 | 0.022056165 | 0.702831 |
| Q7ZVK5 | 0.02218554 | 0.874527 |
| Q6GMJ2 | 0.022757786 | 3.22566 |
| F1QXG4 | 0.022863357 | 2.82874 |
| Q1LWD7 | 0.0229562 | -1.77951 |
| F1QEW4 | 0.023227368 | 0.942526 |
| Q7ZW09 | 0.023480103 | 1.86285 |
| E7FAD0 | 0.023546155 | 3.10134 |
| Q6DGX1 | 0.023694088 | 2.40918 |
| Q7ZVF1 | 0.023799619 | -1.32043 |
| I3ISJ2 | 0.023912776 | 2.01248 |
| E9QEN2 | 0.023915529 | 2.09861 |
| F1QYW8 | 0.02414182 | 2.65206 |
| Q499A7 | 0.024306924 | 2.19476 |
| F1QNB3 | 0.024346134 | 2.09184 |
| Q9YH92 | 0.024444998 | -1.15699 |
| Q6PHU9 | 0.02457084 | 1.88207 |
| Q90XP6 | 0.024592348 | -2.08989 |
| Q7T3F0 | 0.024670048 | 0.779289 |
| Q7T368 | 0.024950543 | 2.30072 |
| E9QFX0 | 0.025318035 | 1.84349 |
| A9JTF6 | 0.025414408 | -1.6273 |
| Z4YIM6 | 0.025920299 | 1.8635 |
| Q6TNV6 | 0.025955536 | 2.17882 |
| Q7SZQ7 | 0.025965698 | -1.54404 |
| Q6NSM6 | 0.026027353 | 1.55449 |
| Q6NYN6 | 0.026145684 | 1.27416 |
| Q642H3 | 0.026485611 | 0.96228 |
| Q9I8U9 | 0.026687015 | 2.90303 |
| Q5NJJ5 | 0.026706686 | -0.206728 |
| A4JYP6 | 0.026821928 | -0.48507 |
| E4VNZ2 | 0.027116273 | 1.80182 |
| F1R6V4 | 0.027384827 | 2.54389 |
| Q6NY49 | 0.02757211 | 1.05739 |
| Q68EH2 | 0.027649036 | 5.08607 |
| Q803P9 | 0.027668142 | 0.779991 |
| A4IG36 | 0.027765788 | -0.842255 |
| Q19U11 | 0.027854156 | -0.980916 |
| B2GQ43 | 0.028578537 | 2.93751 |
| E7F4F3 | 0.028718384 | 0.951628 |
| A8E7N5 | 0.02872632 | -0.926525 |
| X1WEV8 | 0.028730289 | 0.201747 |
| E9QCG8 | 0.028842971 | 2.73333 |
| Q6TH07 | 0.029285996 | 2.80224 |
| Q803X9 | 0.029527046 | -1.34449 |
| Q6IQ92 | 0.02970845 | 3.16474 |
| F1RDA6 | 0.029723504 | 0.594812 |
| E9QH32 | 0.029761855 | -1.05076 |
| Q6PBR5 | 0.029977817 | 2.5901 |
| B8A4B0 | 0.029983339 | 2.64132 |
| F1R9C6 | 0.029987482 | 1.86189 |
| C5NU62 | 0.030424258 | 2.83243 |
| Q6NV46 | 0.030500714 | -1.55084 |
| Q90XR8 | 0.030782224 | -1.64218 |
| Q8AXX5 | 0.03082904 | -0.655121 |
| Q6IQR3 | 0.03084182 | 6.62793 |
| Q64HD0 | 0.031122896 | -1.38414 |
| F1QKX8 | 0.031138666 | -2.3318 |
| H9GX86 | 0.031426788 | 3.78899 |
| E7F9M7 | 0.031436921 | 1.42795 |
| Q802F5 | 0.031762158 | -1.08898 |
| B8JJ32 | 0.032130686 | 2.77611 |
| F1Q670 | 0.032270077 | -1.71662 |
| Q6P948 | 0.032814062 | 1.92839 |
| E9QIF5 | 0.032844299 | -1.7298 |
| Q503F1 | 0.032937452 | 1.95261 |
| F4ZGF2 | 0.033279746 | 1.06184 |
| F1QNI9 | 0.033289709 | 5.02495 |
| Q6PC49 | 0.033319617 | 3.13535 |
| E9QGI1 | 0.033382588 | -0.660227 |
| F1REU6 | 0.03369768 | -1.256 |
| Q5U3U1 | 0.033727178 | -1.22132 |
| Q32PS5 | 0.03390783 | 1.08851 |
| Q6NWL9 | 0.034206607 | 2.41666 |
| Q6P5M2 | 0.034423102 | 0.893567 |
| Q9I8U7 | 0.03456607 | 1.90599 |
| Q6P608 | 0.034692852 | -1.56081 |
| E7EZJ2 | 0.035026763 | -0.331424 |
| E7FAN2 | 0.035205458 | -3.14608 |
| Q6IQ56 | 0.035514851 | -1.0019 |
| Q6Q415 | 0.035547577 | 2.62834 |
| Q7SX99 | 0.036447689 | 3.47002 |
| E7EZ94 | 0.037547411 | 2.37711 |
| Q1LVV8 | 0.037767653 | 3.15441 |
| E7F169 | 0.037893088 | -1.10388 |
| F1R632 | 0.037956836 | 0.873911 |
| Q66I01 | 0.038165416 | 2.08649 |
| Q6IQM8 | 0.038413157 | -1.40476 |
| E9QHN0 | 0.038691004 | 1.85906 |
| B3DJB0 | 0.038933191 | -2.94718 |
| Q9DFT9 | 0.039075091 | -1.97797 |
| F1RDE6 | 0.039149841 | -2.49665 |
| E9QIS9 | 0.039344135 | -1.48227 |
| F1QPE9 | 0.039592233 | -2.10803 |
| F1Q9D1 | 0.039962256 | 1.97065 |
| Q7ZUI0 | 0.04028747 | 2.39498 |
| Q6PBI3 | 0.040757731 | -0.690159 |
| Q1LVG7 | 0.041179399 | 1.92123 |
| E7F971 | 0.041341859 | -1.81237 |
| B0UYN4 | 0.041459112 | 1.27584 |
| Q6NYV3 | 0.04152121 | 3.1052 |
| Q0IJ38 | 0.041944981 | -1.73288 |
| Q66HX0 | 0.042041674 | -2.00434 |
| E7FG54 | 0.042338942 | 1.01492 |
| Q568E8 | 0.042460001 | -0.815934 |
| A2RUX5 | 0.042591212 | -1.61227 |
| Q66IF0 | 0.043224502 | -2.2228 |
| Q6IQS6 | 0.043717971 | 1.20124 |
| F1QKC0 | 0.043841964 | 1.99942 |
| F1Q8J9 | 0.043871249 | 2.63557 |
| Q7ZVY9 | 0.043885394 | 1.25056 |
| E9QEE4 | 0.044420148 | 1.22915 |
| F1Q9Q1 | 0.04459642 | -0.458567 |
| Q2MJQ8 | 0.044687906 | 1.87331 |
| A8DZD1 | 0.044706431 | -2.13322 |
| Q6PBH6 | 0.045041205 | 2.58726 |
| B0S5J9 | 0.045217859 | 0.77997 |
| F1RAM9 | 0.045252231 | 1.1816 |
| Q66HY6 | 0.045353416 | 0.929653 |
| Q7ZWD8 | 0.045404615 | 0.662427 |
| Q90ZA7 | 0.045496711 | 0.830927 |
| Q4VBU2 | 0.045753045 | 3.83661 |
| Q6AXI7 | 0.045899716 | 0.78285 |
| A2BGE0 | 0.04600023 | -1.11524 |
| Q6NYR9 | 0.046263661 | 1.66557 |
| Q6P6E7 | 0.046277512 | 3.30822 |
| Q90Z10 | 0.047273744 | 2.6708 |
| A4FVJ1 | 0.04769908 | -0.54293 |
| E9QDX8 | 0.047739735 | 1.6545 |
| Q4VBU0 | 0.047975554 | 2.08533 |
| F1QEB6 | 0.048069544 | -0.747078 |
| E9QJK1 | 0.048128242 | 2.27805 |
| Q6PC24 | 0.04859706 | -1.54366 |
| B0JZP4 | 0.048803391 | 2.27965 |
| Q5F0G5 | 0.049125841 | 1.45094 |
| F1QXQ1 | 0.049863183 | 3.01684 |
| Q90X41 | 0.049880408 | -0.877574 |
| Q6NV37 | 0.05007489 | 3.57974 |

**Supplemental Table 3:** Differentiated expressed proteins for the Mixture (101µM) exposed group.

| **Protein IDs** | **P value** | **Fold Change** |
| --- | --- | --- |
| Q1LXP8 | 0.000847793 | -1.57266 |
| A2CJ02 | 0.001000115 | 3.54708 |
| Q5XJ96 | 0.001357 | -1.54172 |
| F1Q6Z7 | 0.00155984 | -3.04002 |
| Q6P0V6 | 0.002316701 | -0.89198 |
| F6P1Y9 | 0.002319583 | 0.775496 |
| A8DZ95 | 0.002846754 | -2.2342 |
| Q90XR8 | 0.003270996 | -0.685425 |
| A8DZD1 | 0.003535086 | 1.38809 |
| E7F5V5 | 0.004056113 | -1.19324 |
| Q567N9 | 0.004627112 | -0.531658 |
| Q4QRD2 | 0.004724219 | 1.01152 |
| F1QHK3 | 0.006201975 | -1.40435 |
| Q499A7 | 0.006355065 | -2.55822 |
| Q801U1 | 0.006562661 | -1.38913 |
| Q6DBX3 | 0.007254041 | -2.55597 |
| F1QNX5 | 0.008206727 | 0.856861 |
| F1QD90 | 0.008668221 | 0.257477 |
| Q642H3 | 0.008685604 | 1.26684 |
| F1QJR3 | 0.010123484 | -2.35102 |
| Q6DGX1 | 0.01115347 | -1.92636 |
| E7FAN2 | 0.011371821 | 0.782455 |
| Q1LWN2 | 0.011710084 | 6.22967 |
| F1R3Y6 | 0.012621472 | 1.95769 |
| Q5RGI5 | 0.01388769 | -0.161459 |
| F8W450 | 0.01507544 | -0.346007 |
| F1Q4Q4 | 0.01572389 | 1.60257 |
| F1Q7H4 | 0.016083479 | -0.671977 |
| A9JRE2 | 0.016341052 | -0.53529 |
| Q1L8A0 | 0.017536786 | 0.227477 |
| Q1LVG4 | 0.018472259 | -0.455594 |
| Q8UVW9 | 0.018541292 | -0.270947 |
| F1QLA4 | 0.018608441 | 1.10141 |
| Q1JPZ7 | 0.018657781 | -0.207977 |
| Q6DEJ9 | 0.0196074 | 1.0243 |
| E7F1Y1 | 0.020131216 | -1.11419 |
| F1QZL8 | 0.020760127 | 0.96029 |
| A3KPW8 | 0.021113003 | 0.472631 |
| Q7T028 | 0.02148523 | -0.492602 |
| E4VNZ2 | 0.02214522 | 0.337053 |
| B3DI91 | 0.023110003 | -2.50447 |
| Q9DGL3 | 0.023994958 | -0.663114 |
| F1QH84 | 0.024688233 | 1.07547 |
| X1WHD9 | 0.02469733 | -1.74574 |
| Q498X8 | 0.024941927 | 0.373582 |
| E9QC40 | 0.025901803 | 0.763173 |
| A8E5C5 | 0.026500862 | -1.58908 |
| F1R3X2 | 0.027742781 | -0.450203 |
| A2BGE0 | 0.028014315 | -0.434714 |
| F1QQY8 | 0.028513465 | -1.16585 |
| F1RBG8 | 0.029164867 | -1.32753 |
| F8W2B6 | 0.029463206 | 1.95208 |
| H0WF22 | 0.029464563 | 2.18607 |
| A2AR65 | 0.029689986 | -2.16504 |
| Q1MTC4 | 0.030235698 | 4.70664 |
| Q5RHV1 | 0.032848081 | -0.623653 |
| Q9I8V1 | 0.032968562 | -0.906849 |
| Q6NY41 | 0.033409502 | 0.787936 |
| Z4YIQ5 | 0.033861797 | 0.738782 |
| E7F9B4 | 0.033963309 | 0.486539 |
| F1R074 | 0.034568457 | -0.794447 |
| X1WFW6 | 0.035631163 | -1.7661 |
| E7F4F3 | 0.035953403 | -2.29401 |
| A8KBQ5 | 0.036700333 | 1.0654 |
| G1K2X0 | 0.038596685 | -0.341554 |
| A3KPI6 | 0.038847225 | -0.850215 |
| Q5SP90 | 0.040030406 | 0.935036 |
| F1R161 | 0.040431514 | -1.21081 |
| Q2MJQ8 | 0.041338051 | 1.05129 |
| Q7ZTW5 | 0.04189093 | 0.678208 |
| Q7ZVD2 | 0.041928565 | 0.490545 |
| A4JYP8 | 0.042176448 | -0.812987 |
| E7F2E9 | 0.044695109 | -0.728691 |
| F1QWM3 | 0.047203043 | 2.34129 |
| F1QPG1 | 0.047332561 | 1.82221 |
| Q7ZV03 | 0.047621163 | 1.30953 |
| E7EXU3 | 0.048104976 | 2.03211 |
| Q7T368 | 0.048706845 | -0.922369 |
| E7F2L6 | 0.049701558 | 1.06424 |

**Supplemental Table 4:** Differentiated expressed proteins for the BMAA (100µM) exposed group.

| **Canonical Pathways** | **BMAA**  **(100µM)** | **MCLR**  **(1µM)** | **Mixture**  **(101µM)** |
| --- | --- | --- | --- |
| RhoGDI Signaling | N/A | -2 | -2.121 |
| Signaling by Rho Family GTPases | N/A | 2 | 2.121 |
| ILK Signaling | N/A | N/A | 2.646 |
| Calcium Signaling | N/A | N/A | -2.449 |
| Hepatic Fibrosis Signaling Pathway | N/A | N/A | 2.121 |
| D-myo-inositol-5-phosphate Metabolism | N/A | N/A | -2 |
| 3-phosphoinositide Biosynthesis | N/A | N/A | -2 |
| Apelin Cardiomyocyte Signaling Pathway | N/A | N/A | 2 |
| Regulation of Actin-based Motility by Rho | N/A | N/A | 1.89 |
| RhoA Signaling | N/A | N/A | 1.89 |
| Superpathway of Inositol Phosphate Compounds | N/A | N/A | -1.342 |
| Gα12/13 Signaling | N/A | N/A | 1.342 |
| Phospholipase C Signaling | N/A | N/A | 1.342 |
| 3-phosphoinositide Degradation | N/A | N/A | -1 |
| Netrin Signaling | N/A | N/A | -1 |
| Systemic Lupus Erythematosus In B Cell Signaling Pathway | N/A | N/A | -1 |
| Systemic Lupus Erythematosus In T Cell Signaling Pathway | N/A | N/A | -0.816 |
| Cardiac Hypertrophy Signaling | N/A | N/A | 0.816 |

**Supplemental Table 5:** List of all canonical pathway activated or inhibited under each condition and their respective z scores.

| **Symbol** | **Gene Name** | **UniProt Accession** | **Expr Fold Change** | **Expr p-value** | **Location** |
| --- | --- | --- | --- | --- | --- |
| MYBPC3 | myosin binding protein C, cardiac | F1Q615 | 4.28158 | 0.014810851 | Cytoplasm |
| CDH17 | cadherin 17 | F1RD80 | 3.32832 | 0.003288365 | Plasma Membrane |
| MYBPH | myosin binding protein H | Q6GMJ2 | 3.22566 | 0.022757786 | Cytoplasm |
| MYH4 | myosin heavy chain 4 | E7FAD0 | 3.10134 | 0.023546155 | Cytoplasm |
| TNNT3 | troponin T3, fast skeletal type | Q9I8U9 | 2.90303 | 0.026687015 | Cytoplasm |
| MYL4 | myosin light chain 4 | Q4QRD2 | 2.77618 | 0.013451168 | Cytoplasm |
| SYN1 | synapsin I | B0UXI4 | 2.1441 | 0.004944814 | Plasma Membrane |
| MYLPF | myosin light chain, phosphorylatable, fast skeletal muscle | E9QG51 | 1.91464 | 0.000440677 | Cytoplasm |
| MYL1 | myosin light chain 1 | Q9I8U7 | 1.90599 | 0.03456607 | Cytoplasm |
| Cdc42 | cell division cycle 42 | Q52WY2 | 1.85253 | 0.004992522 | Plasma Membrane |
| CTSB | cathepsin B | E9QDX8 | 1.6545 | 0.047739735 | Cytoplasm |
| MYOC | myocilin | Q5F0G5 | 1.45094 | 0.049125841 | Cytoplasm |
| MYL2 | myosin light chain 2 | E7FG54 | 1.01492 | 0.042338942 | Cytoplasm |
| LCT | lactase | F1QBK3 | -1.42172 | 0.015897177 | Plasma Membrane |
| MYBPC2 | myosin binding protein C, fast type | Q0IJ38 | -1.73288 | 0.041944981 | Cytoplasm |
| SLC8A1 | solute carrier family 8 member A1 | B3DJB0 | -2.94718 | 0.038933191 | Plasma Membrane |
| ARFGEF1 | ADP ribosylation factor guanine nucleotide exchange factor 1 | E7FGL2 | -3.05072 | 0.007318296 | Cytoplasm |
| ANXA1 | annexin A1 | Q804H2 | 4.10424 | 0.007339391 | Plasma Membrane |
| HSPD1 | heat shock protein family D (Hsp60) member 1 | Q803B0 | 3.58377 | 0.019529 | Cytoplasm |
| NDUFA13 | NADH:ubiquinone oxidoreductase subunit A13 | Q6PC49 | 3.13535 | 0.033319617 | Cytoplasm |
| GLUD1 | glutamate dehydrogenase 1 | B8A4B0 | 2.64132 | 0.029983339 | Cytoplasm |
| UQCRC1 | ubiquinol-cytochrome c reductase core protein 1 | Q6PBH6 | 2.58726 | 0.045041205 | Cytoplasm |
| TARDBP | TAR DNA binding protein | Q802C7 | 2.55077 | 0.015899007 | Nucleus |
| UQCRQ | ubiquinol-cytochrome c reductase complex III subunit VII | Q6DGN1 | 2.43111 | 0.015047003 | Cytoplasm |
| SUCLG1 | succinate-CoA ligase alpha subunit | Q6DGX1 | 2.40918 | 0.023694088 | Cytoplasm |
| ATP5MF | ATP synthase membrane subunit f | Q6PBH7 | 2.26218 | 0.005512137 | Cytoplasm |
| PPIA | peptidylprolyl isomerase A | Q499A7 | 2.19476 | 0.024306924 | Cytoplasm |
| MSI2 | musashi RNA binding protein 2 | X1WC29 | 2.13454 | 0.009539596 | Cytoplasm |
| BCKDHA | branched chain keto acid dehydrogenase E1 subunit alpha | Q4VBU0 | 2.08533 | 0.047975554 | Cytoplasm |
| UQCRH | ubiquinol-cytochrome c reductase hinge protein | I3ISJ2 | 2.01248 | 0.023912776 | Cytoplasm |
| PYCR2 | pyrroline-5-carboxylate reductase 2 | Q503F1 | 1.95261 | 0.032937452 | Cytoplasm |
| NDUFA6 | NADH:ubiquinone oxidoreductase subunit A6 | Q8AW03 | 1.94919 | 0.014109122 | Cytoplasm |
| ATP5F1D | ATP synthase F1 subunit delta | Q1LVG7 | 1.92123 | 0.041179399 | Cytoplasm |
| PHGDH | phosphoglycerate dehydrogenase | Q6PHU9 | 1.88207 | 0.02457084 | Cytoplasm |
| KPNA4 | karyopherin subunit alpha 4 | Q7ZVY9 | 1.25056 | 0.043885394 | Nucleus |
| GRPEL1 | GrpE like 1, mitochondrial | Q32PS5 | 1.08851 | 0.03390783 | Cytoplasm |
| RDH13 | retinol dehydrogenase 13 | Q6NY49 | 1.05739 | 0.02757211 | Cytoplasm |
| NME4 | NME/NM23 nucleoside diphosphate kinase 4 | E9QH32 | -1.05076 | 0.029761855 | Cytoplasm |
| SLC16A10 | solute carrier family 16 member 10 | A1L1W9 | -1.08241 | 0.01196961 | Plasma Membrane |
| PPP3CA | protein phosphatase 3 catalytic subunit alpha | E7F169 | -1.10388 | 0.037893088 | Cytoplasm |
| STAT3 | signal transducer and activator of transcription 3 | Q6NV46 | -1.55084 | 0.030500714 | Nucleus |
| PRSS27 | serine protease 27 | F1QS02 | -1.99283 | 0.018042637 | Extracellular Space |
| CFB | complement factor B | F1R886 | -2.13874 | 0.009098923 | Extracellular Space |
| AMPH | amphiphysin | Q6P960 | -2.21665 | 0.0081313 | Plasma Membrane |

**Supplemental Table 6:** List of all DEPs associated with interactive network for skeletal/muscular disorder and cellular assembly/organization.
