## Supplementary figures and images for "Exposure to a mixture of BMAA and MCLR synergistically modulates behavior in larval zebrafish while exacerbating molecular changes related to neurodegeneration"

### Supplemental Figure 1

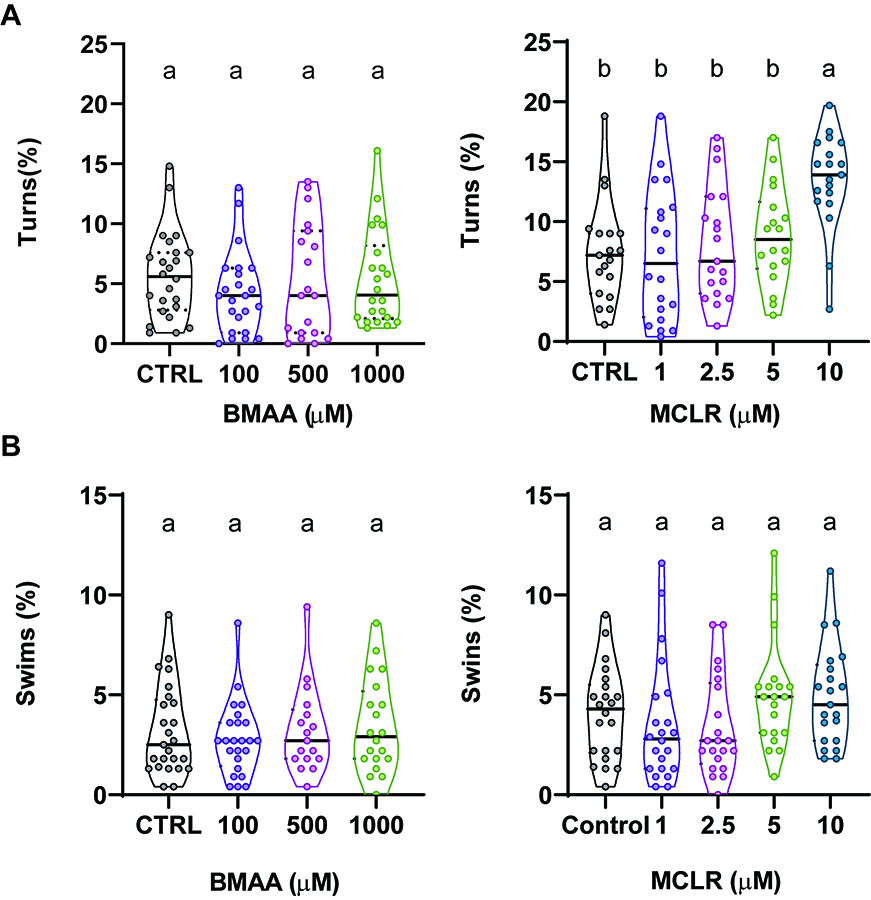

### Supplemental Figure 2

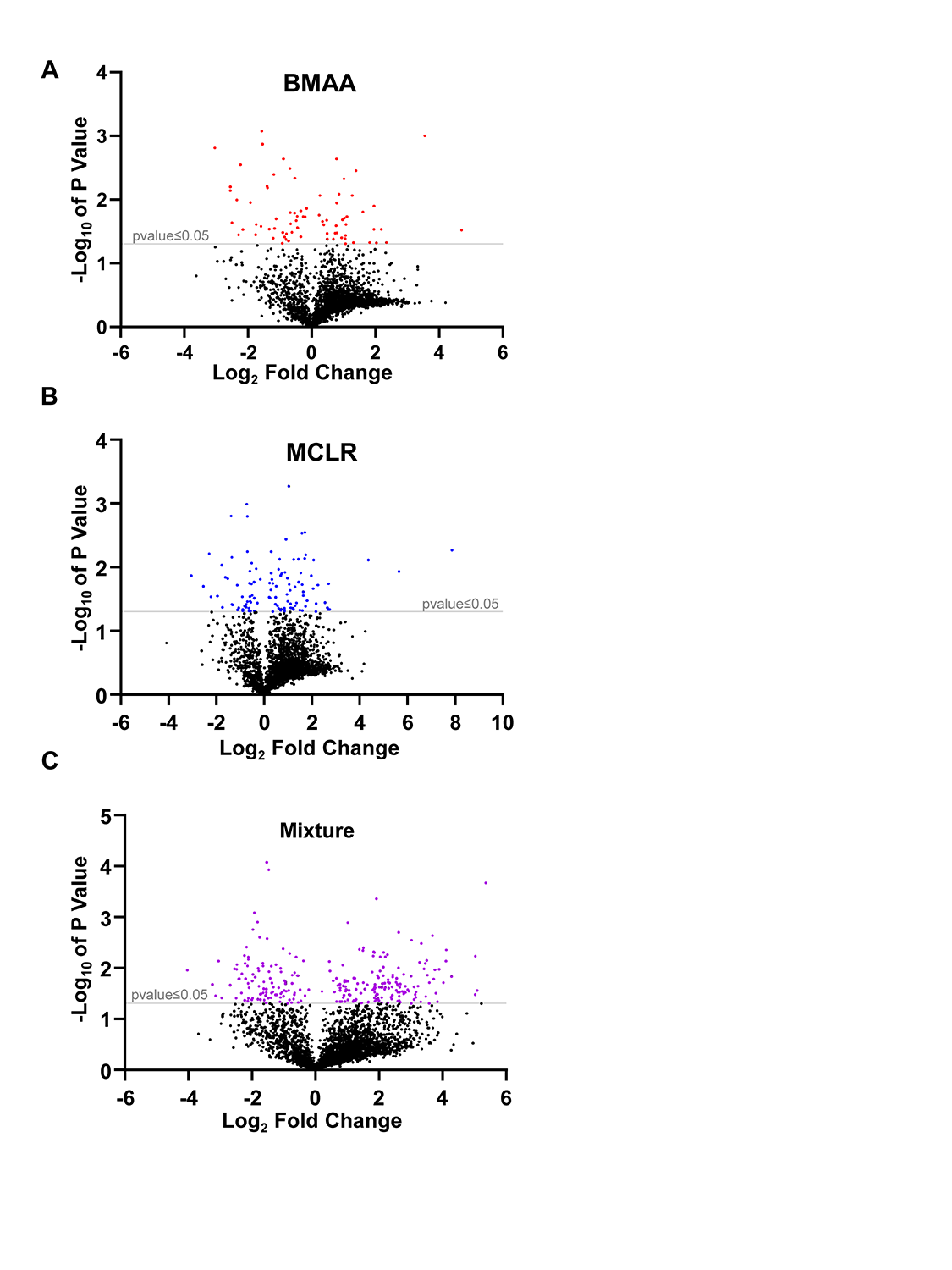
